# A *Toxoplasma gondii* putative arginine transporter localizes to the plant-like vacuolar compartment and controls parasite extracellular survival and stage differentiation

**DOI:** 10.1101/2023.08.31.555807

**Authors:** Federica Piro, Silvia Masci, Geetha Kannan, Riccardo Focaia, Tracey L. Schultz, Vern B. Carruthers, Manlio Di Cristina

## Abstract

*Toxoplasma gondii* is a protozoan parasite that infects a broad spectrum of hosts and can colonize many organs and cell types. The ability to reside within a wide range of different niches requires substantial adaptability to diverse microenvironments. Very little is known about how this parasite senses various milieus and adapts its metabolism to survive, replicate during the acute stage, and then differentiate to the chronic stage. Most eukaryotes, from yeast to mammals, rely on a nutrient sensing machinery involving the TORC complex as master regulator of cell growth and cell cycle progression. The lysosome functions as a signaling hub where TORC complex assembles and is activated by transceptors, which both sense and transport amino acids, including the arginine transceptor SLC38A9. While most of the TORC components are lost in *T. gondii*, indicating the evolution of a distinct nutrient sensing mechanism, the parasite’s lysosomal plant-like vacuolar compartment (PLVAC) may still serve as a sensory platform for controlling parasite growth and differentiation. Using SLC38A9 to query the *T. gondii* proteome, we identified four putative amino acid transporters, termed TgAAT1-4, that structurally resemble the SLC38A9 arginine transceptor. Assessing their expression and sub-cellular localization, we found that one of them, TgAAT1, localized to the PLVAC and is necessary for normal parasite extracellular survival and bradyzoite differentiation. Moreover, we show that TgAAT1 is involved in the PLVAC efflux of arginine, an amino acid playing a key role in *T. gondii* differentiation, further supporting the hypothesis that TgAAT1 might play a role in nutrient sensing.

**IMPORTANCE:** *T. gondii* is a highly successful parasite infecting a broad range of warm-blood organisms including about one third of all humans. Although *Toxoplasma* infections rarely result in symptomatic disease in individuals with a healthy immune system, the incredibly high number of persons infected along with the risk of severe infection in immunocompromised patients and the potential link of chronic infection to mental disorders make this infection a significant public health concern. As a result, there is a pressing need for new treatment approaches that are both effective and well-tolerated. The limitations in understanding how *Toxoplasma gondii* manages its metabolism to adapt to changing environments and triggers its transformation into bradyzoites have hindered the discovery of vulnerabilities in its metabolic pathways or nutrient acquisition mechanisms to identify new therapeutic targets. In this work, we have shown that the lysosome-like organelle PLVAC, acting through the putative arginine transporter TgAAT1, plays a pivotal role in regulating the parasite’s extracellular survival and differentiation into bradyzoites.

## Introduction

Cell growth is a process that requires a precise and tight regulation in both unicellular and multicellular organisms of all kingdoms. Prokaryotes have a more direct and simple response to nutrient availability to regulate their growth while eukaryotes, particularly multicellular organisms, have evolved a more complicated network that controls cell growth and division. TORC1 and 2 are the master regulators of cell growth, proliferation, differentiation, metabolism, and survival in most eukaryotes from yeast to mammals (1–4). Lysosomes function as a signaling hub where TORC1 complex assembles and is activated by nutrient sensors such as the arginine transceptor SLC38A9 (5–7). SLC38A9 is an eleven trans-membrane protein that localizes on lysosome membrane and senses lysosomal luminal arginine to activate TORC1 through its cytosolic N-terminal region of 119 amino acids that interacts with the Rag-Ragulator complex in a vacuolar ATPase (V-ATPase) dependent manner (8–11). Specifically, TORC1 activation by SLC38A9, together with Ragulator, seems to rely on its function as a non-canonical GEF for Rag GTPase (12, 13). Furthermore, recent studies have also shown that SLSC38A9 facilitates the efflux of several essential amino acids from the lysosome in an arginine-dependent fashion, especially leucine (14). Additionally, lysine promotes the interaction of SLC38A9 and the Rag-Ragulator complex, suggesting that lysine could also activate TORC1 through this arginine sensor (15). Notably, other amino acid transporters such as the heterodimeric transporter SLC7A5 (LAT1)/SLC3A2, recruited to the lysosomal membrane by LAPTM4b (16, 17), and PAT1 (SLC36A1) have been shown to stimulate the activity of TORC1 in response to other amino acids (18–21). It is now accepted that the role of amino acid transporters in amino acid sensing is not only due to their ability to translocate amino acids through the lysosome membrane, but also to work as activators of amino-acid dependent signaling independently of an effective transport. With a dual receptor-transporter function, these lysosomal amino acid transporters have been termed ‘transceptors’ for their roles in sensing amino acid availability upstream of intracellular signaling pathways (22, 23).

Despite a comprehensive understanding of the mechanisms that regulate cell growth in most eukaryotes, these processes are still poorly understood in apicomplexan parasites. Even though the evolutionary age of the TOR pathway indicates that both TORC1 and TORC2 were present in the last eukaryotic common ancestor (LECA), the majority of TORC1 pathway components have been lost or significantly modified in the apicomplexan lineage (24–27). How these organisms detect and respond to nutrient changes is still elusive. As obligate intracellular parasites, they face unique challenges for nutrient acquisition and regulation due to their dependency on host cell resources. The loss of TORC1 pathway components in the apicomplexan lineage suggests that these organisms have undergone adaptations to suit their parasitic lifestyle and nutrient acquisition strategies. In fact, *Plasmodium* does not have the two well-characterized mechanisms to sense amino acid fluctuations, the TORC complex and the GCN2-downstream transcription factors (27). However, malaria parasites appears to encode a downstream component of the TORC pathway, the putative ortholog of the RNA polymerase (Pol) III repressor Maf1, which has been shown to modulate pol III transcription in a TOR-dependent manner in other eukaryotes (28). PfMaf1 plays an important role in maintaining parasite viability during starvation and other forms of stress. Furthermore, in a recent work, the Mota lab has identified 3 distinct parasite kinases, Nek4, eIK1 and eIK2, that function as key regulators of parasite growth under amino acid scarcity (29). This study showed that malaria parasites can activate a stage specific kinase in response to a decrease in methionine and isoleucine abundance that ultimately leads to a reprograming of replication and parasite development.

As for *T. gondii*, knowledge on mechanisms regulating its growth and differentiation is much more limited than in *Plasmodium*. *T. gondii* tachyzoites survive for only a short time extracellularly (30–33), and they replicate exclusively inside host cells, like most apicomplexan life stages (34). *T. gondii* expresses two GCN2-like eIF2a kinases that play key roles in the parasite adaptation to changes in the environment (32, 35). One of these kinases, TgIF2K-C, is required to promote survival of intracellular tachyzoites cultured under glutamine starvation conditions (35) while the second one, TgIF2K-D, promotes survival of tachyzoites upon egress from host cells (32). Regarding the TORC complex, akin malaria parasites, *T. gondii* seems to have lost the majority of components (25–27). A putative Tor homologue is present in *T. gondii* genome (36), TGME49_316430, but the fitness score of this gene is positive (37), indicative of its dispensability. Deletion of this gene showed no apparent phenotype in *in vitro* experiments (Di Cristina et al. in preparation) in line with its positive fitness score. Furthermore, this gene encodes a protein whose size is nearly twice that of yeast and mammal Tor, suggesting that may have evolved a different role in a context where all the other TORC components are lost. However, even if a different nutrient sensing machinery has evolved in *T. gondii*, the lysosome may have maintained its role of central signaling hub where nutrient sensors regulate parasite growth, differentiation, and response to environmental changes. Moreover, arginine, the TORC1 activator amino acid sensed by the lysosomal transporter SLC38A9, is a key determinant of *T. gondii* differentiation, since its starvation was found to trigger differentiation of replicative tachyzoites into bradyzoites (38). *T. gondii* possesses a lysosome-like organelle termed the plant-like vacuolar compartment (PLVAC) that plays multiple functions as digestive organelle, being involved in ion storage and homeostasis, endocytosis, and autophagy (39). So far, only one PLVAC transporter has been characterized, the *T. gondii* homologue of *Plasmodium* chloroquine resistance transporter (TgCRT), which likely exports products of proteolytic digestion (peptides, amino acids) from the PLVAC into the parasite cytoplasm (40).

The aim of the work presented herein was to identify additional PLVAC amino acid transporters and assess whether they play any role in parasite survival, growth, or differentiation. Here, we have identified bioinformatically 4 putative amino acid transporters that we named TgAAT1-4. One of them, TgAAT1, localizes to the PLVAC, likely exports arginine from PLVAC to the parasite cytosol, and is necessary for normal extracellular survival and stable inter-conversion between tachyzoites to bradyzoites in response of differentiation stimuli.

## Results

### Identification of SLC38A9 homologues in *T. gondii*

The lysosomal amino acid transporter SLC38A9 is a component of the lysosomal sensing machinery that signals amino acid availability to TORC1 (9). We wondered whether *Toxoplasma* similarly exploits amino acid transporters to regulate its growth and/or differentiation. To address this, we interrogated the *Toxoplasma* genome database ToxoDB (41) using the human SLC38A9 as query to identify putative *T. gondii* SLC38A9 homologues. This search returned three hits that displayed modest homology to the query but are annotated as amino acid transporters (Fig. S1). The three genes encoding these proteins (TGME49_227430, TGME49_227570, and TGME49_227580) are all located on chromosome X. TGME49_227570 and TGME49_227580 are adjacent to one another in a head-to-tail arrangement, whereas TGME49_227430 is ∼30 kbp away and in the opposite orientation. The similarity and close linkage of these genes suggests they arose by gene duplication from a common progenitor. A fourth hit displaying lower homology with SLC38A9 (Fig. S2), TGME49_226060, was also annotated as amino acid transporter and is likewise located on chromosome X, but ∼2 million base pairs away from the other three putative genes. TGME49_226060 also displayed moderate similarity (19.0% identity, Fig. S3) to the *Plasmodium falciparum* amino acid transporter PfAAT1, but showed good superimposition of their Alphafold2 predicted 3D structures (Fig. S4). Due to this homology, the protein encoded by TGME49_226060 was named TgAAT1, and TGME49_227430, TGME49_227570, and TGME49_227580, were named TgAAT2-4, respectively. Despite low amino acid conservation with SLC38A9, structural analysis of the TgAAT proteins using Alphafold2, TOPCONS, and I-TASSER software suggested the presence of 11 transmembrane domains that can be aligned by ClustalW with those of SLC38A9 (Fig. 1). The DALI server identified SLC38A9 as the best 3D superimposed structure for all the four TgAAT proteins (Fig. 2 shows 3D superimposition of SCL38A9 and TgAAT1 as representative example). Together, these findings suggest that *T. gondii* possess four putative amino acid transporters with structural similarity to SLC38A9, which prompted us to characterize these integral membrane proteins further.

**Figure 1.**
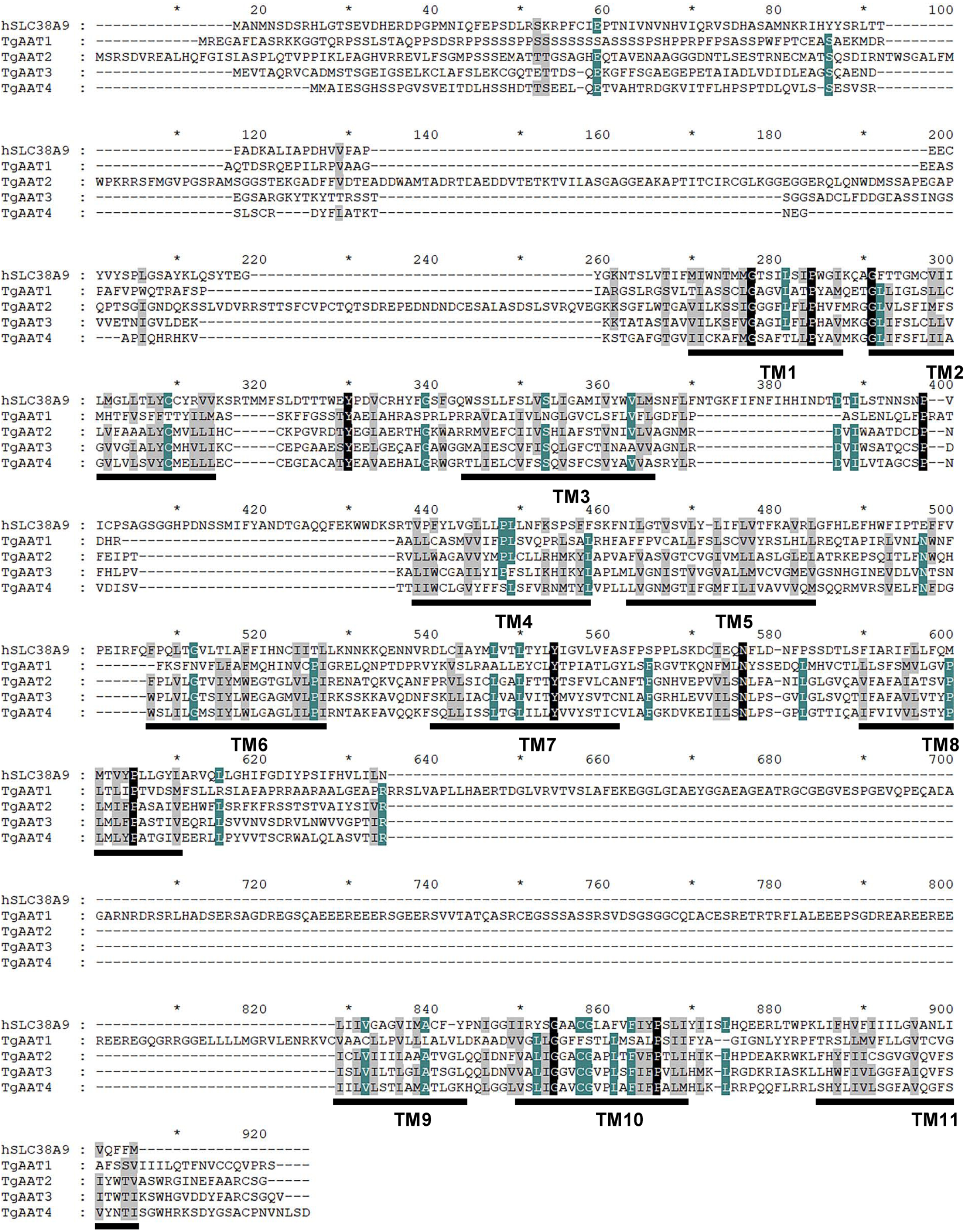
Multiple sequence alignment of SLC38A9 and putative amino acid transporters (TgAATs) using ClustalW software. Predicted transmembrane domains are underlined and number as TM1-11. Accession number of proteins used in the alignment analysis are as follows: SLC38A9, homo sapiens, accession number NP_001336312.1; ToxoDB accession number for TgAATs: TgAAT1, TGME49_226060; TgAAT2, TGME49_227430; TgAAT3, TGME49_227570; TgAAT4, TGME49_227580.

**Figure 2.**
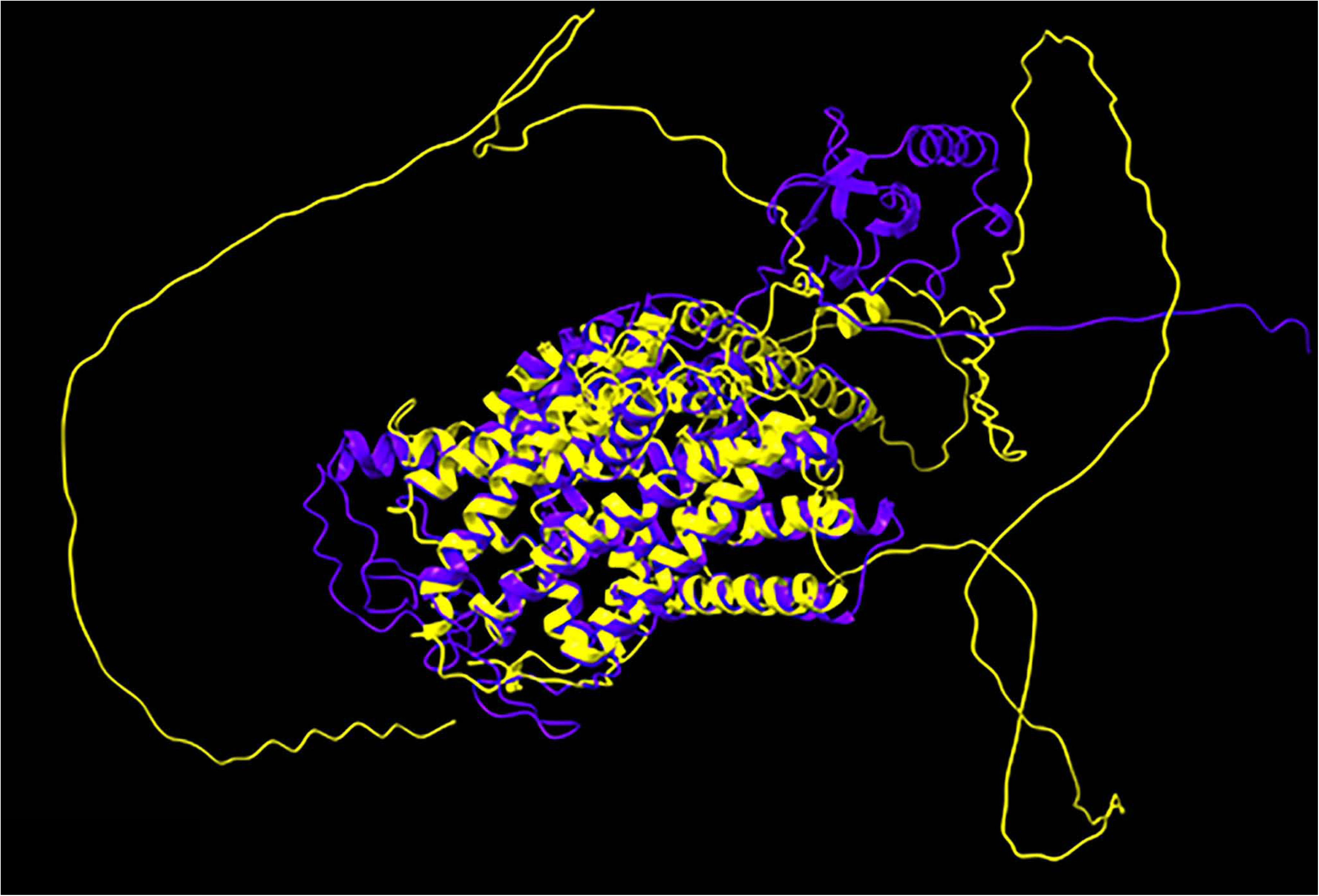
AlphaFold2 model and structural comparison of SLC38A (Blue) and TgAAT1 (Yellow).

### TgAAT1 is a resident protein of the *T. gondii* PLVAC

To determine the expression profile of the four TgAAT genes, we performed a series of semiquantitative RT-PCR experiments using RNA extracted from type II PruΔ*ku80/Sluc* strain (42)(hereafter called Pru) tachyzoites or *in vitro* differentiated bradyzoites. TgAAT2 and TgAAT4 were expressed at similar levels in both stages, while TgAAT1 showed low expression in tachyzoites and higher expression in bradyzoites (Fig. 3). TgAAT3 was not detected in Pru tachyzoites or bradyzoites but was expressed in type I RHΔ*ku80*Δ*hxprt* strain tachyzoites (Fig. 3B). Based on their expression in Pru, we endogenously tagged TgAAT1, TgAA2, and TgAAT4 with 4 copies of the c-myc epitope (4x c-myc) at their N- or C-terminal ends (Fig. S5). Disappointing, none of the tagged strains was positive by immunofluorescence (IFA) or western blotting (WB) (data not shown). Failure to detect the four proteins could be explained by either low expression or cleavage of tags, particularly for the C-terminus with its predicted orientation toward the hydrolytic lumen of the PLVAC. To account for both potential explanations, we generated vectors to transiently overexpress each protein with a single N-terminal c-myc epitope. For TgAAT2 and TgAAT4, we again failed to detect expression by IFA or WB, possibly due to a tight regulation of translation, poor protein stability, or loss of the c-myc epitope due to N-terminal processing. However, transient overexpression of TgAAT1 showed that it occupies the PLVAC based on colocalization with the PLVAC marker cathepsin protease L (TgCPL) (Fig. S6A and B). Furthermore, WB analysis of tachyzoites transiently transfected with c-mycTgAAT1 showed a band of the expected molecular weight of ⁓75 kDa (Fig. S6C). Since over-expression of TgAAT1 resulted in an appreciable signal by both IFA and WB, we generated a new strain in Pru where the *TgAAT1* promoter was replaced with that of *TgSAG1* together with adding a c-myc epitope at the N-terminus end of TgAAT1 (Fig. S6D-F). The higher expression resulted from the stronger *TgSAG1* promoter confirmed the PLVAC localization of TgAAT1 by IFA (Fig. S6G). We also corroborated the PLVAC localization of TgAAT1 by endogenous inserting a c-myc epitope (Fig. S7) into a large loop between the eighth and the ninth transmembrane domains oriented towards the cytosolic face of the PLVAC (Fig. 4A and B). TgAAT1 staining in this latter strain was also detected in *in vitro* cysts after one week of differentiation induced by alkaline media (Fig.4C). Collectively, these findings indicate that TgAAT1 localizes to the PLVAC, which prompted us to focus our investigation on it for the remainder of the study.

**Figure 3.**
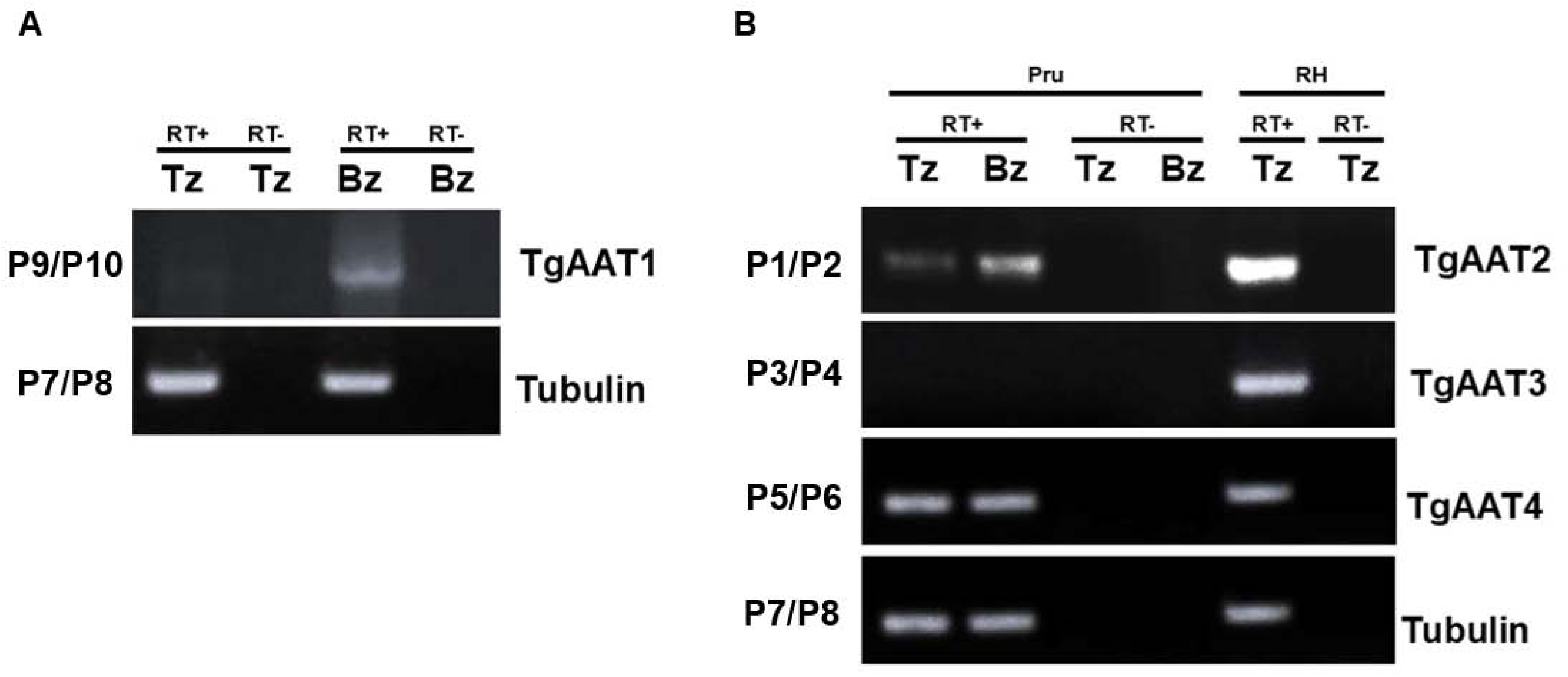
Semiquantitative RT-PCR analysis of TgAATs expression in either the tachyzoite or bradyzoite stages. RNA isolated from *in vitro* tachyzoites (Tz) or bradyzoites (Bz) was reverse transcribed and used for PCR analysis using primers P9/P10 for TgAAT1 (panel A), P1/P2 for TgAAT2 (top panel B), P3/P4 for TgAAT3 (middle panel B) and P5/P6 for TgAAT4 (bottom panel B). Tubulin was amplified using primers P7/P8 (last of both panel A and B). Positive (‘RT+’) and negative (‘RT–’) signs indicate RNAs with or without reverse transcriptase, respectively. The representative agarose gel electrophoresis of RT-PCR products showed the expected bands of sizes: 100, 380, 400, 500 and 250 bp for TgAAT1, 2, 3, 4 and tubulin, respectively.

**Figure 4.**
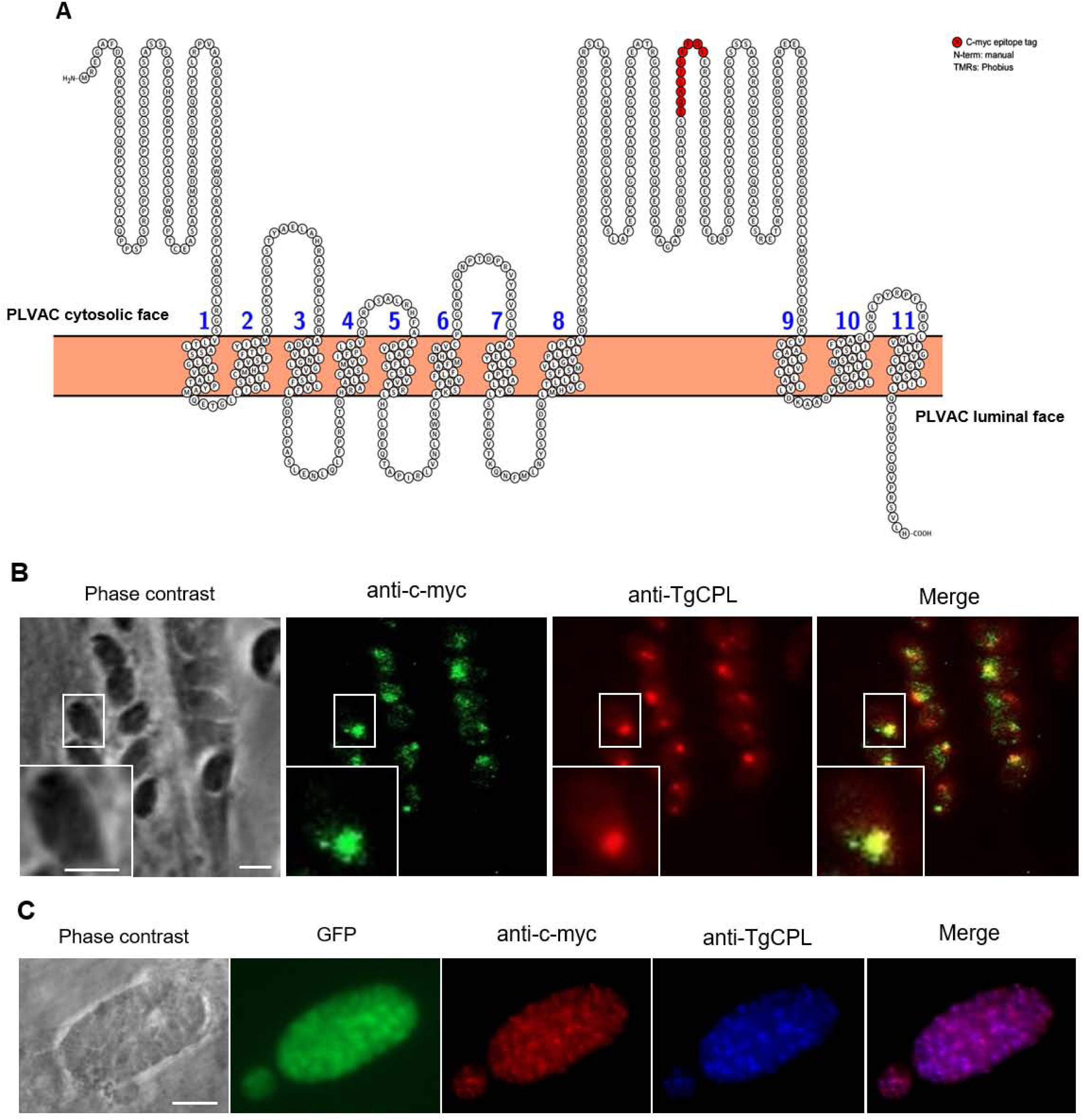
Analysis of TgAAT1 sub-cellular localization indicates that the protein is resident on the PLVAC. A) TgAAT1 was internally tagged with the c-myc epitope. Protter analysis (https://wlab.ethz.ch/protter/<u>#</u>) of TgAAt1 amino acid sequence showing the integration of the c-myc tag in the loop between the predicted 8^th^ and 9^th^ transmembrane domains. B) Intracellular TgAAT1 tachyzoites were stained with anti-c-myc (green) and anti-TgCPL (red) to mark the PLVAC (scale bar 10µm). TgAAT1 and TgCPL co-localization indicated that TgAAT1 is resident on the PLVAC (inset, scale bar 2µm). C) TgAAT1 staining (red) of *in vitro* bradyzoites overlapped with that of TgCPL (blue), confirming the localization on PLVAC (scale bar 20µm). EGFP is expressed in bradyzoites of all strains derived from the parental PruΔ*ku80/Sluc*.

### TgAAT1 plays a key role in extracellular tachyzoite survival

To investigate the role of the TgAAT1 protein in the life cycle of *T. gondii*, we used CRISPR/Cas9 technology (43) to generate a TgAAT1 knock-out strain, PΔ*aat1* (Fig. S8A and B). A complement strain was also generated by re-expressing TgAAT1 as a cDNA driven by its own promoter and integrated in the tubulin locus (PΔ*aat1*:AAT1, Fig. S8C and D). Characterization of these strains using classical assays such as plaque, invasion, replication, and competition assays allowed the identification of a strong defect in extracellular survival of PΔ*aat1*, that was rescued in the complement strain. Extracellular survival of the KO strain was impaired within 0.5 h of host cell extrusion, even if parasites were in a rich media (DMEM) and was markedly reduced to 5-30% after 2 h of extracellular incubation (Fig. 5A). The shorter tolerance to extracellular life of parasites lacking TgAAT1 may be responsible of the reduced number of plaques observed in the plaque assay (Fig. 5B). The parental strain also outcompeting PΔ*aat1* in a co-infection assay (Fig. 5C). More specifically, co-infecting host cells with parental and PΔ*aat1* strains in a 1:4 ratio at passage 0 (P0) resulted in the inversion of this ratio to ∼4:1 after P5, whereas no significant competition was seen between the parental and complement strains. Replication assays and intracellular tachyzoite viability showed no significant differences among the three strains (Fig. 5D and E), in line with similar plaque sizes (data not shown). Invasion assays revealed a modest phenotype, perhaps due to the reduced extracellular survival of PΔ*aat1* tachyzoites (Fig. 5F). Together these findings indicate that TgAAT1 promotes extracellular survival of *T. gondii* tachyzoites.

**Figure 5.**
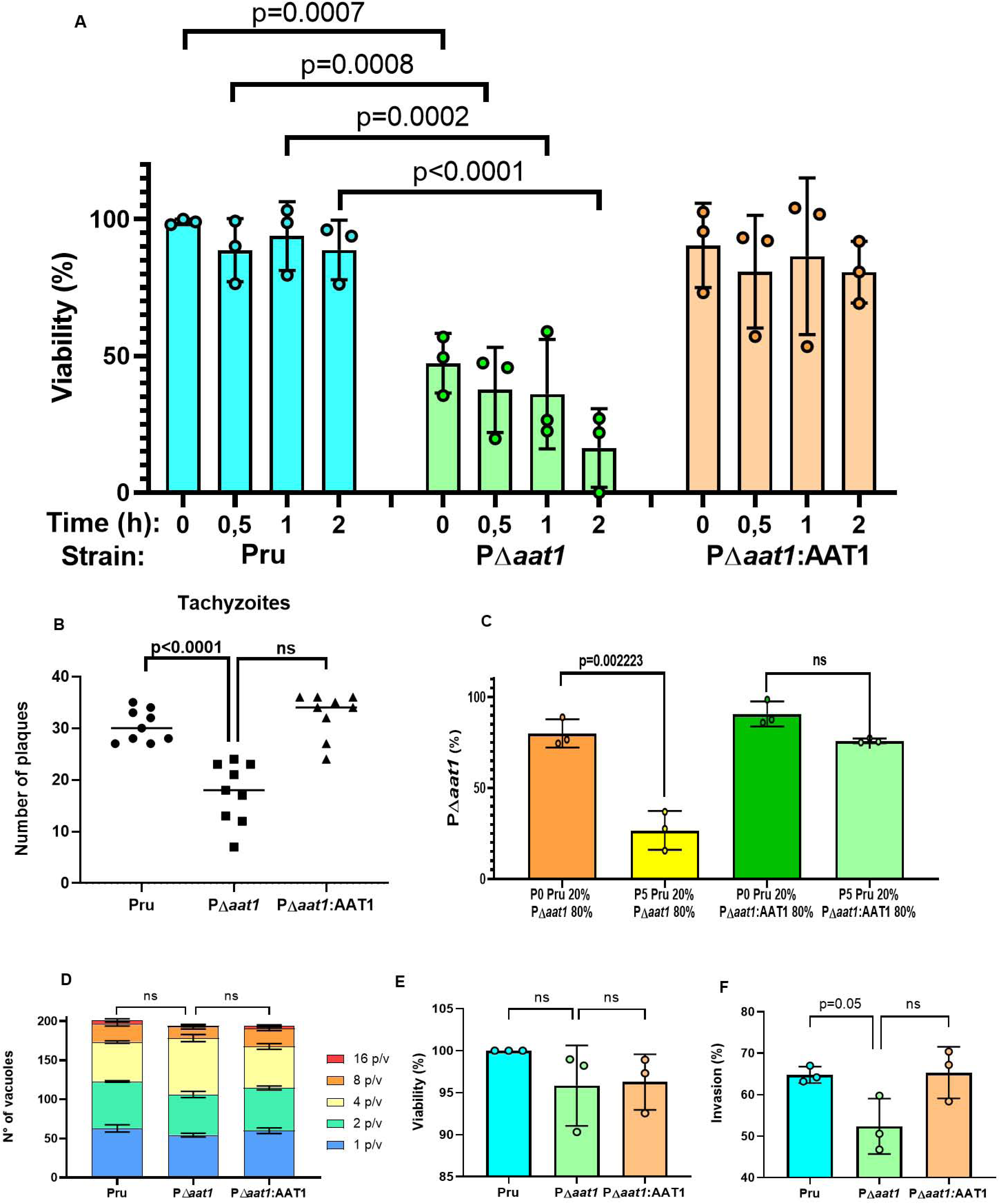
Phenotypic analysis of TgAAT1 tachyzoites. A) TgAAT1 promotes survival of tachyzoites upon egress from host cells. Tachyzoites of Pru, PΔ*aat1* and PΔ*aat1*:AAT1 were mechanically liberated from HFF cells and 10^6^ parasites of each strains were incubated for 0, 0.5, 1 and 2 h in D10 media at 37°C and 5% CO_2_. After extracellular incubation, tachyzoite viability was assessed by PMA/qPCR analysis. One-way ANOVA with Holm-Sidak’s multiple comparisons was used to compare the medians of data combined from 3 biological replicates. B) Plaque assay of Pru, PΔ*aat1* and PΔ*aat1*:AAT1 tachyzoites showed that deletion of TgAAT1 resulted in a reduced number of plaques compared to both the parental and complement strain. Unpaired student’s t-test was used to compare the medians of data combined from 3 biological replicates. C) Competition assay to compare the fitness of the PΔ*aat1* and complement tachyzoites with that of the Pru strain. HFF monolayers grown in T25 flasks were infected with 10^6^ tachyzoites of a mixture of either parental and PΔ*aat1* or parental and PΔ*aat1*:AAT1 strains in the ratio of 1:4. Parasites were grown for five passages and then harvested, purified and used for qPCR analysis to calculate the ratio between the two mixed strains. The presence of the DHFR selection cassette in the PΔ*aat1* and PΔ*aat1*:AAT1 strains but not in the parental was exploited to selectively amplify only the gDNA of these strains using primers binding to the tubulin promoter and DHFR cDNA of the selection cassette (see Table S1). Primers binding to the tubulin gene were used to normalize gDNA quantity since they amplified the gDNA of all three strains. The results showed the inversion of the 1:4 ratio in 4:1 after 5 passages of the only mixture parental/ PΔ*aat1*. The graph represents means +/- the SD of three independent experiments. Unpaired student’s t-test was performed. D) TgAAT1 ablation did not caused any replication defect. Parasites were cultured in monolayer HFF cells and samples were collected at 24 h post-infection, fixed, stained with DAPI (4′,6-diamidino-2-phenylindole) and anti-TgSAG1, and quantified by fluorescence microscopy. At least 100 vacuoles were counted from 6 different fields of view. The percentages of different replication stages in the population for each strain were plotted. Results represent means ± SD from three independent experiments. Two-way ANOVA with Dunnett’s multiple comparisons was performed. p/v: parasites/vacuole. E) PMA/qPCR viability assay of intracellular Pru, PΔ*aat1* and PΔ*aat1*:AAT1 tachyzoites. TgAAT1 ablation did not impact on intracellular tachyzoite viability. The graph represents means +/- the SD of three independent experiments. One-way ANOVA with Holm-Sidak’s multiple comparisons was performed. F) TgAAT1 ablation slightly impacts on host cell invasion. Shown are the results of a red-green invasion assay of tachyzoites after 20 min of incubation with HFF cells. Parasites were stained as described in Materials and Methods. The graph represents means +/- the SD of three independent experiments. One-way ANOVA with Holm-Sidak’s multiple comparisons was performed.

### TgAAT1 deficient bradyzoites are less viable *in vitro*, but show increased cyst burden in chronically infected mice

To determine whether ablation of TgAAT1 affected cyst formation, morphology, or viability, we analyzed the phenotype of PΔ*aat1* during the chronic stage of *T. gondii in vitro* and *in vivo*. Analysis of *in vitro* cysts showed that ablation of TgAAT1 caused altered morphology and formation of translucent vacuoles and small dark dots in bradyzoites (Fig. 6A). Although the nature of these alterations was not further investigated, PΔ*aat1* bradyzoites were ⁓40-50% less viable than parental and complement strains after 1 or 2 weeks of *in vitro* conversion (Fig. 6B and C). Unexpectedly, CBA/J mice infected with 10^5^ PΔ*aat1* tachyzoites showed an ⁓10-fold increase in brain cyst burden 5 weeks post-infection compared to those inoculated with the parental or complement strains (Fig. 6D). This *in vivo* analysis was repeated three times and each experiment consisted of five mice per group infected with the same strain. Together these findings suggest that although *in vitro* bradyzoites lacking TgAAT1 show altered morphology and viability, they conversely produced more brain cysts in infected mice.

**Figure 6.**
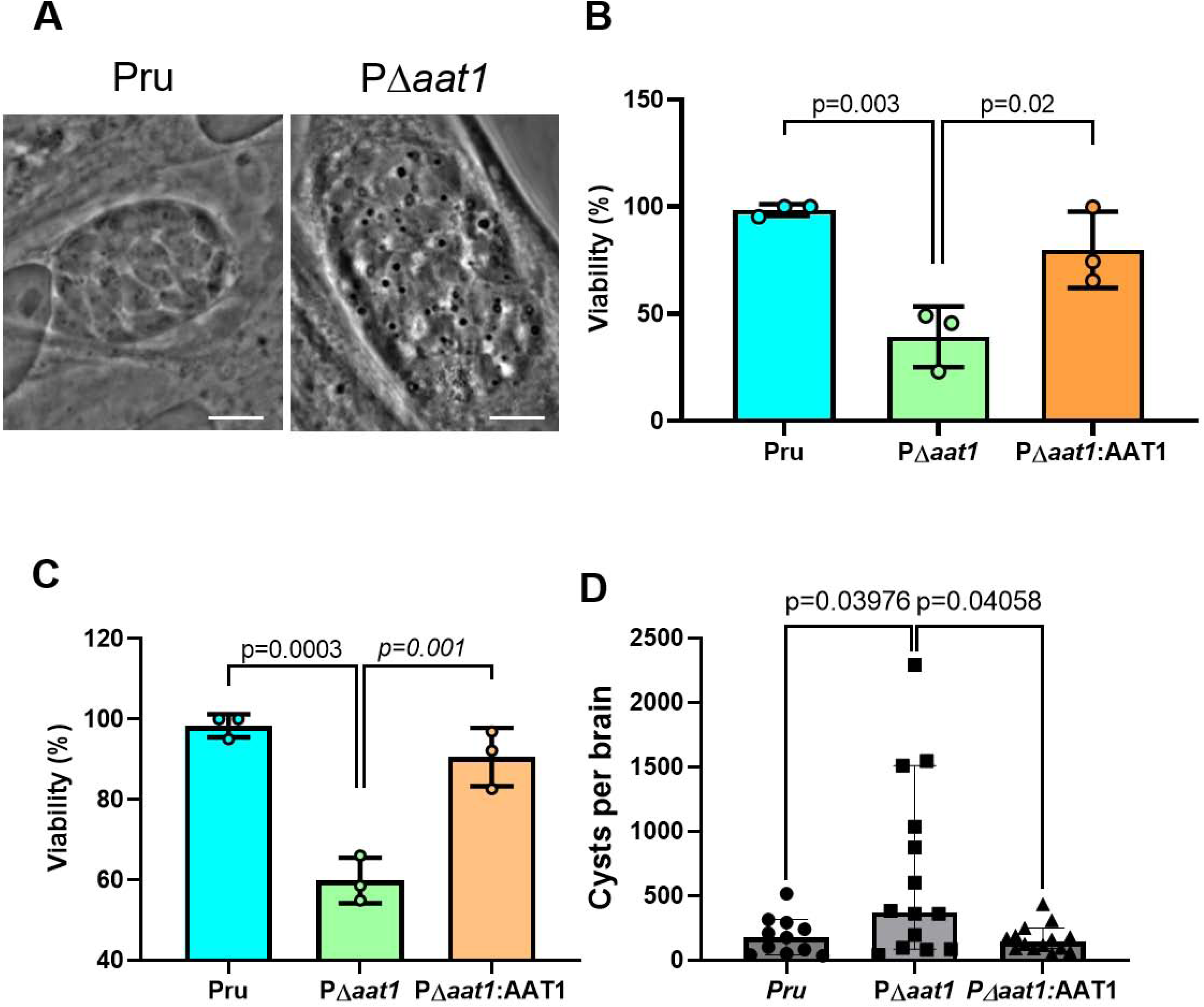
Deletion of the TgAAT1 gene led to significant changes in the formation, viability and morphology of cysts containing bradyzoites. A) Representative images of parental or PΔ*aat1 in vitro* cyst containing bradyzoites. Parasites were grown in alkaline media for 2 weeks and then cysts were analyzed using optical microscopy under a 100X objective. TgAAT1 ablation caused alteration of morphology and formation of translucent vacuoles and small dark dots in bradyzoites (scale bar = 10µm). B-C) TgAAT1-deficient bradyzoites displayed reduced viability. Pru, PΔ*aat1* and PΔ*aat1*:AAT1 tachyzoites were converted into bradyzoites by alkaline induction for either 1 week (B) or 2 weeks (C) and analyzed for viability using the PMA/qPCR assay. Error bars indicate the S.D. from three independent experiments. One-way ANOVA with Holm-Sidak’s multiple comparisons was performed. D) Three groups composed of five CBA/J mice were infected with either the wild-type (parental), PΔ*aat1* or PΔ*aat1*:AAT1 strain. The experiment was repeated three times, resulting in a total of 15 mice for each group in the study. The graph depicts the number of cysts present in the brains of survival mice from each group 5 weeks post-infection. The PΔ*aat1* strain exhibited a higher number of cysts per brain compared to mice infected with both the wild-type and complement strain. This suggests that TgAAT1 ablation might have an impact on the parasite’s ability to form cysts within the host. Mann Whitney test was performed.

### PΔ*aat1* showed unstable inter-conversion between tachyzoites and bradyzoites

Brain cyst burden in infected mice is likely influenced by several factors including the timing of tachyzoite-bradyzoite conversion and strength of cues that the parasite uses for such conversion *in vivo*. The increased cyst burden observed in PΔ*aat1* infected mice prompted us to assess the rate and efficiency of tachyzoite to bradyzoite inter-conversion. To this end, we measured the fluorescence intensity of parasite-containing vacuoles stained for either the tachyzoite marker TgSAG1 or the bradyzoite markers EGFP and TgBAG1 over the first 10 days of *in vitro* differentiation in alkaline medium. EGFP, which is driven by the LDH2 promoter, is an early bradyzoite marker, whereas TgBAG1 expression increases later during differentiation. Parental and complement strains showed the typical rate of conversion characterized by a gradual decrease in TgSAG1 during the first 4 days in alkaline medium, followed by the nearly complete loss of TgSAG1 and the appearance of the EGFP and TgBAG1 from days 5 and 7 onwards, respectively (left and right panels of Fig. 7). In contrast, the same analysis showed that ablation of TgAAT1 destabilized the inter-conversion between tachyzoites and bradyzoites (middle panels of Fig. 7). Notably, the tachyzoite marker TgSAG1 persisted at higher level during the entire period of differentiation including an apparent further increase at day 10 post alkaline induction. Both early and late bradyzoite markers, EGFP and TgBAG1, displayed an up and down expression with a drop of their signal intensity around days 9-10 post induction, coinciding with the TgSAG1 peak. TgBAG1 in PΔ*aat1* parasites failed to reach the same intensity of signals observed in the parental and complement strains at any point during the 10 days in alkaline medium. Together these findings suggest that PΔ*aat1* parasites show an unstable commitment to bradyzoite conversion under *in vitro* differentiation conditions.

**Figure 7.**
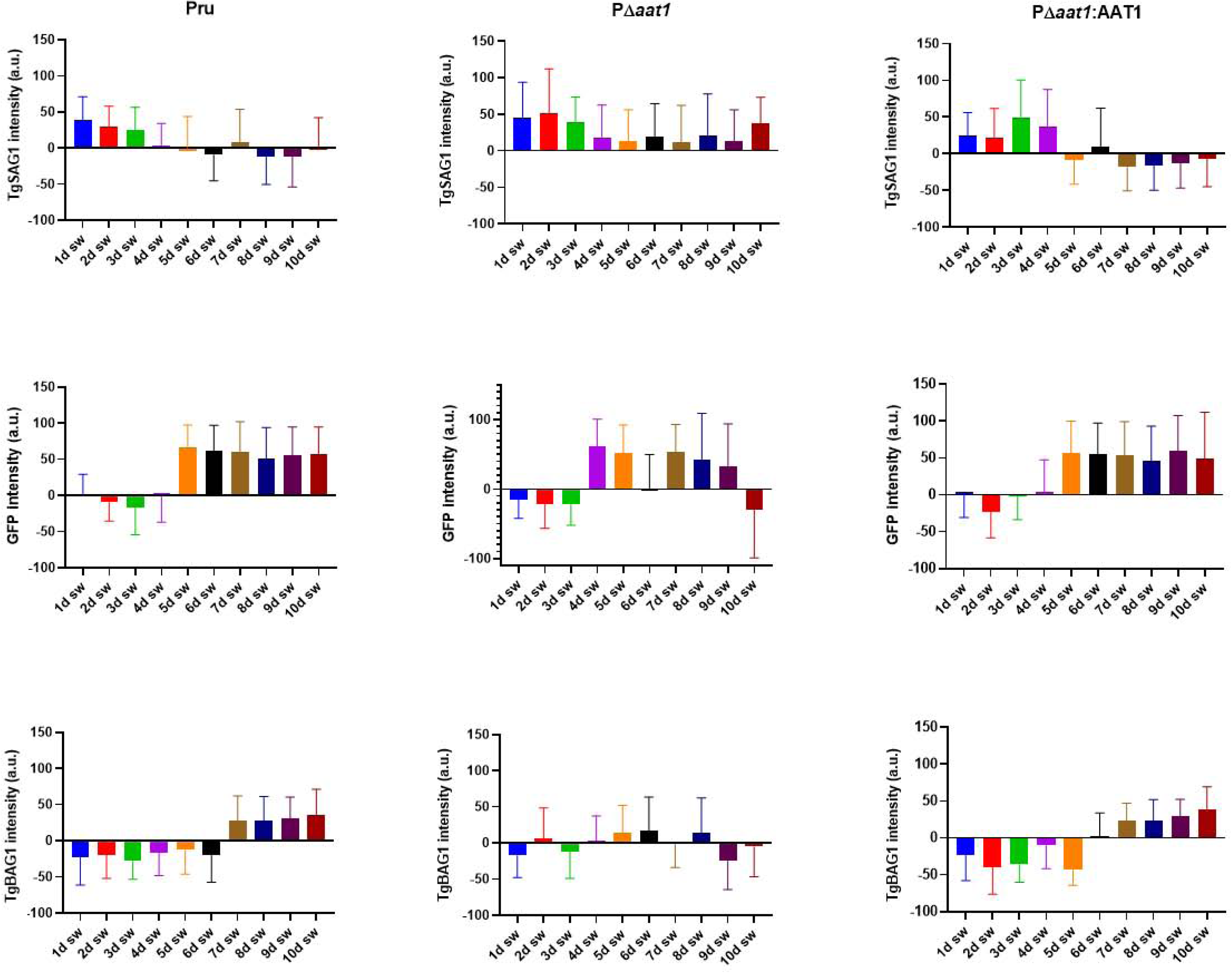
Ablation of TgAAT1 gene leads to alterations in the tachyzoite-bradyzoite inter-conversion process. Bradyzoite conversion kinetics of Pru, PΔ*aat1* or PΔ *aat1*:*AAT1* strains were assessed in *in vitro* alkaline differentiation. Three groups of 20 HFF monolayers grown on coverslip were infected with 10^5^ tachyzoites of each strain. Tachyzoites were induced to convert into bradyzoites 24 h post-infection by replacing the D10 media with alkaline media. Two samples from each group were fixed everyday from day 1 to day 10 post-conversion and stained with either rabbit anti-TgSAG1 or rabbit anti-TgBAG1 followed by anti-rabbit Alexa 594. Intensity of fluorescence for TgSAG1, TgBAG1, and EGFP was measured in 300 vacuoles from each slide by fluorescence microscopy. The results showed that deletion of the TgAAT1 gene caused changes in the behavior of the parasites, possibly affecting their ability to maintain the bradyzoite stage.

### TgAAT1 is involved in arginine efflux from the PLVAC

TgAAT1 is annotated as an amino acid transporter and its 3D structure prediction and comparison with other proteins support this notion. Structural comparison of TgAAT1 using DALI server identified the arginine transporter SLC38A9 as its best 3D superimposition hit. To investigate whether TgAAT1 was also capable of transporting arginine or other amino acids that are sensed to regulate cell proliferation, we used an esterified amino acid approach similar to that employed by Verdon et al. (44). Esterified amino acids are permeable to membranes and thus enter any subcellular compartment including lysosomes. Lysosomal acid hydrolases remove the methyl-ester group, thereby causing the accumulation of the corresponding amino acid inside the lysosome, ultimately generating an osmotic effect (45). Cells can alleviate this stress by using an amino acid transporter(s) to efflux substrate amino acid(s) from the lysosome to the cytosol. Deleting a transporter can result in slower or absent efflux from the lysosome of the accumulated amino acid(s) that may become toxic for the cell that eventually dies. We selected 6 amino acids: arginine, glutamine, leucine, lysine, because they are involved in TORC1 activation in other organisms (46, 47), and tryptophan and glycine as representative amino acids not involved in lysosomal nutrient sensing. Intracellular tachyzoites of the parental, TgAAT1 KO, and complement strains were mechanically extruded from HFFs, filter purified, and incubated for 2 h at 37°C in either normal RPMI media or RPMI supplemented with one of the six selected esterified amino acid. The concentration of each esterified amino acid used in the assay was the highest tolerated with no toxicity to the parental strain. After incubation, the viability of parasites was measured by propidium monoazide–quantitative polymerase chain reaction (PMA-qPCR) assay adapted from other studies (48–57). As shown in figure 8, treatment with esterified arginine caused a ⁓60% decrease of parasite viability exclusively in the PΔ*aat1* strain, whereas the other 5 esterified amino acids showed no significative effects in any of the parental, knockout, or complement strains. This result suggested that the lack of TgAAT1 limited efflux of arginine from the PLVAC, thereby reducing the capacity of KO parasites to rapidly compensate the osmotic effect generated by over-loading the organelle with this amino acid. To further test this possibility, we developed another assay also based on the esterified amino acid but with a different readout. Scerra et al. (58) showed that reduced efflux of an amino acid from the lysosomes because of the lack of its transporter resulted in its accumulation within the organelle and consequent osmotic stress that caused the enlargement of the lysosomes. Thus, we applied a similar protocol to test whether any of the six selected amino acids caused the enlargement of the PLVAC of parasites lacking TgAAT1. To this end, we first endotagged the PLVAC protein TgFYVE (59) at its N-terminus with mCherry in the 3 strains (Fig. S9) and performed IFA using anti-mCherry antibodies to stain the PLVAC membrane where this protein localized (Fig. 9A). Exploiting this staining, we measured the area of 900 PLVACs of parental, PΔ*aat1*, and complement tachyzoites. This analysis showed that arginine was again the only amino acid of the six selected amino acids that caused a significative enlargement of the PLVAC in PΔ*aat1* compared to the other 2 strains (Fig. 9B). Collectively, these assays provide indirect evidence that TgAAT1 is necessary for normal efflux of arginine from the PLVAC.

**Figure 8.**
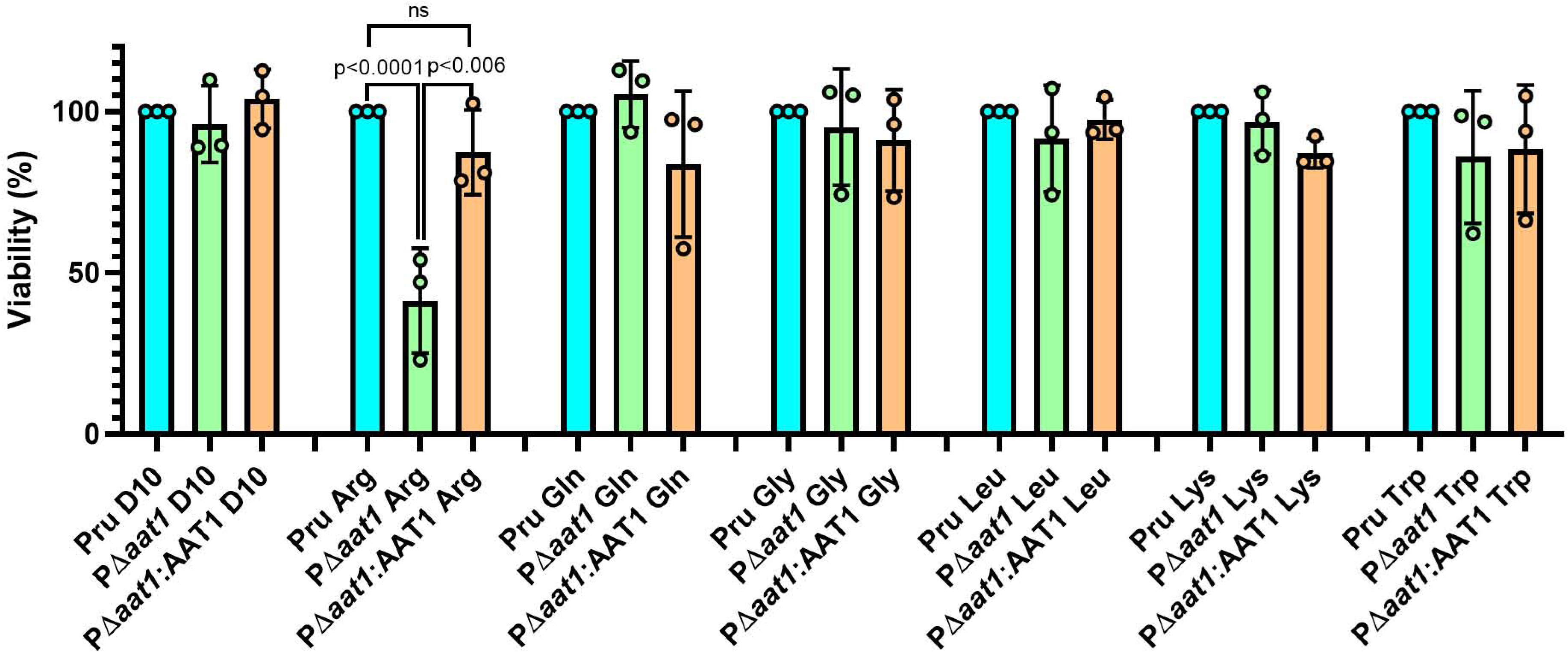
TgAAT1 is involved in arginine efflux from the PLVAC. Extracellular Pru, PΔ*aat1* and PΔ*aat1*:AAT1 tachyzoites were incubated with normal RPMI media or RPMI media supplemented with one of the six different esterified amino acids for 1h at 37°C and analyzed for viability using the PMA/qPCR method. Parasites were incubated with the following esterified amino acid concentrations: 25mM Arg, 100mM Gln, 5mM Gly, 10mM Leu, 10mM Lys,10mM Trp. Error bars indicate the S.D. from three independent experiments. One-way ANOVA with Dunnett’s multiple comparisons was performed.

**Figure 9.**
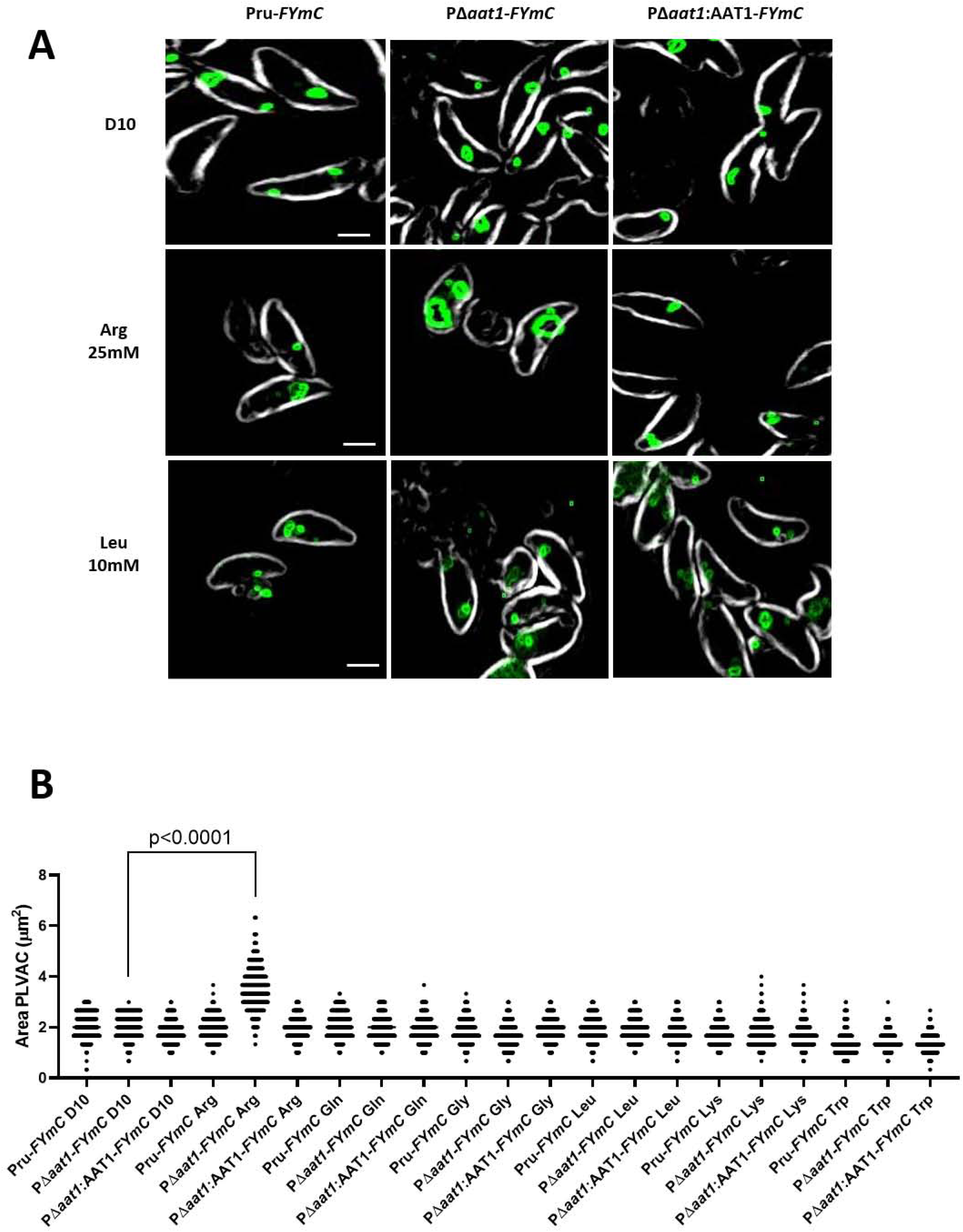
Arginine caused the enlargement of PLVAC in parasites lacking the TgAAT1 gene. A) representative images of Pru*-FYmC*, PΔ*aat1-FYmC* and PΔ*aat1:AAT1-FYmC* tachyzoites incubated with either D10, esterified arginine or esterified leucine for 1 h and stained with anti-mCherry and anti-rabbit Alexa594 (scale bar = 2µm). B) Area of the PLVAC of Pru*-FYmC*, PΔ*aat1*-FYmC and PΔ*aat1:AAT1-FYmC* tachyzoites incubated with either D10 or one of the six esterified amino acids for 1 h. Parasites were incubated with the following esterified amino acid concentrations: 25mM Arg, 100mM Gln, 5mM Gly, 10mM Leu, 10mM Lys,10mM Trp. The area of 300 PLVACs were measured for each sample. One-way ANOVA with Holm-Sidak’s multiple comparisons was performed. Data are presented as mean ± s.d. of three biological replicates, each with 300 PLVACs evaluated.

## Discussion

Among apicomplexans *T. gondii* is an exceptionally adaptable and successful parasite. Its ability to infect a wide variety of hosts, ranging from mammals to birds, is a testament to its versatility. Furthermore, this parasite has evolved strategies that allow it to colonize various organs and cell types within its many hosts, which is a key factor in its ability to persist and propagate. Thus, the adaptability of *T. gondii* is crucial for its survival in all these diverse microenvironments. However, very little is known on how *T. gondii* senses and adapts to different environments, particularly regarding nutrient availability and how this affects its survival, growth, and ability to differentiate between different life stages. Differently from most eukaryotes, *T. gondii* and other apicomplexans do not possess a conserved TORC machinery that regulates their metabolism. Although a gene encoding a potential Tor kinase homologue can be found in ToxoDB (36), so far there is no evidence that this protein is the central regulator of the parasite metabolism (Di Cristina et al., in preparation). Furthermore, homologues of other core components of the nutrient sensing machinery (including Gator, Raptor, Ragulator, Rags, and Lst8) were not identified in *T. gondii* and *Plasmodium spp.*, suggesting that the TORC complex was lost in the Apicomplexa lineage over the course of evolution since it was already present in LECA (24–27). However, even if the TORC complex was lost in these organisms, the lysosome might have retained the function as a metabolic signaling hub for a distinct nutrient sensing machinery (5, 60). *T. gondii* possesses a lysosomal-like organelle named PLVAC (39) whose proteome is still poorly characterized with only a few membrane proteins identified and only one peptide/amino acid transporter, TgCRT (40). Amino acid transporters have been implicated in sensing of specific amino acids from inside the lysosome and modulating activity of the TORC1 complex (61). SLC38A9, a sodium-coupled amino acid transporter, is considered the main lysosome membrane protein that senses lysosomal luminal arginine and activates TORC1 by directly interacting with the Rag-Ragulator complex (11, 62). Such studies of lysosome-based amino acid sensing in other eukaryotes prompted us to address whether the *T. gondii* lysosome-like organelle PLVAC is also involved in regulating any aspect of parasite growth and/or differentiation. TgCRT was shown to be involved in the PLVAC homeostasis, but parasite growth and differentiation were not altered upon ablating TgCRT (63, 64). We then performed searches in ToxoDB using human lysosomal or yeast amino acid vacuolar transporters (avt) to identify additional putative PLVAC amino acid transporters. Results from these analyses consistently identified the same four proteins encoded by TGME49_226060 (TgAAT1), TGME49_227430 (TgAAT2), TGME49_227570 (TgAAT3), and TGME49_227580 (TgAAT4), although amino acid identity was modest and restricted to relatively small regions of these proteins. More convincing, structural analysis of all four proteins using DALI server identified SLC38A9 as best 3D structural homolog. However, despite N- and C-terminal tagging, we never detected signals for TgAAT2, TgAAT3, or TgAAT4 by IFA or WB, not even when overexpressed in transient transfection experiments. Differently, TgAAT1 was clearly expressed in parasites transiently transfected with a vector carrying the coding cDNA under control of the tubulin promoter and fused at the 5’-end with the c-myc sequence. TgAAT1 localized to the PLVAC as its staining in IFA overlapped with that of the organelle marker TgCPL. However, endogenous tagging of the TgAAT1 N-terminus was not detectable unless the promoter was swapped with that of TgSAG1 to increase expression. Although IFA detection of endogenous internally tagged TgAAT1 was weak, it was sufficient to confirm the PLVAC localization in non-overexpressing parasites. Homology searches using TgAAT1 as the query identified orthologues in several apicomplexans including *Plasmodium falciparum*, which expresses PfAAT1 (65). Recent work on PfAAT1 has revealed that localizes to the digestive vacuole (DV) and that mutations of critical residues conferred parasite resistance to chloroquine, similarly to PfCRT (66, 67). Despite the important role played by PfAAT1 in drug resistance, its transporter specificity has not been addressed so far.

Our work on TgAAT1 has allowed the identification of its role in promoting extracellular survival of *T. gondii* tachyzoites. Deletion of TgAAT1 markedly reduced tachyzoite tolerance to extracellular life with decreased viability seen within 0.5 h of mechanical cellular extrusion. This phenotype may explain results obtained from the competition assay where parental parasites outgrew PΔ*aat1*, but not the complement strain, when mixed and allowed to replicate in the same host cell monolayers. Whether TgAAT1 supports extracellular survival directly e.g., by providing amino acids from PLVAC proteolysis, or indirectly e.g., by working as a sensor that activates a survival pathway remains to be determined and deserves further investigation. Differently, viability of intracellular tachyzoites was not affected by the lack of TgAAT1. Interesting, *in vitro* PΔ*aat1* bradyzoites were ⁓50% less viable than parental and complement strains. Moreover, *in vitro* differentiated PΔ*aat1* bradyzoites showed an altered morphology and the presence of translucent vacuoles and small dark dots within their cytoplasm. Such morphologic changes are reminiscent of those seen in bradyzoites lacking TgCPL. Similarly, TgCPL proteolytic activity is not critical for tachyzoites but is more important for bradyzoite viability (42). However, cyst burden of mice infected with PΔ*aat1* was 10-fold higher than parental and complement strains. Although this finding was in apparent contradiction with the reduced bradyzoite viability of PruΔ*aat1*, analyses of the rate and efficiency of tachyzoite to bradyzoite inter-conversion over ten days indicated that the lack of PΔ*aat1* caused an unstable differentiation, as we observed many bradyzoites converting back to tachyzoites. This stage instability may be responsible for the higher brain cyst burden observed in mice infected by PΔ*aat1*. Hypothetically, incomplete bradyzoite differentiation, with some of them converting back into tachyzoites, might be responsible for new waves of tachyzoite expansion in the brain before forming new cysts. Also, *T. gondii* might receive stronger cues to differentiate *in vivo*, thus potentially accounting the eventual formation of more cysts in infected mice.

Furthermore, consistent with a putative role of amino acid transporter, we have applied an indirect but previously validated approach based on the use of esterified amino acid precursors (44, 58) to assess whether TgAAT1 can transfer any of the amino acids that are sensed by the nutrient sensing machinery in other eukaryotes. We focused this analysis on the 6 amino acids, arginine, glutamine, leucine, lysine, that are TORC1 activators in other organisms along with tryptophan and glycine as representative amino acids not involved in the amino acid sensing pathway. Notably, arginine was the only amino acid that caused parasite death and PLVAC swelling.

Taken together, these data suggest that TgAAT1 plays a key role in regulating parasite metabolism and differentiation since its ablation results in weaker extracellular tolerance and instability in maintaining the correct parasite form. In one scenario, TgAAT1 might provide amino acids to temporarily sustain *T. gondii* metabolism when parasites lack access to host-derived nutrients. Alternatively, TgAAT1 may work as sensor that regulates parasite metabolism to respond to environment changes. Although we have no direct evidence that TgAAT1 is a nutrient sensor, that TgAAT1 is necessary for normal and stable differentiation between tachyzoites and bradyzoites is indicative of this function. Differently from SLC38A9 (68), this *T. gondii* putative transporter does not possess the long N-terminal tail facing the cytosolic side of the lysosome that interacts with the Rag-Ragulator complex to activate mTORC1, which nevertheless is not found in *T. gondii*. However, TgAAT1 is likely responsible for efflux of arginine through the PLVAC membrane, the amino acid sensed by the transactivator SLC38A9 and that regulates cell growth in higher eukaryotes. Moreover, arginine is known to be a key amino acid regulating *T. gondii* differentiation since growth of tachyzoites in media lacking arginine induced bradyzoite conversion (38).

In conclusion, our work represents a step toward identifying components of nutrient sensing machinery that may control the life cycle of *T. gondii*. Whether TgAAT1 is a nutrient sensor or an indirect player is still to be demonstrated but our findings again show the central role played by the PLVAC in an aspect of parasite metabolism.

## MATERIALS AND METHODS

### Host cell and parasite cultures

Human foreskin fibroblasts (HFFs) were grown and maintained in Dulbecco’s modified Eagle’s medium containing 10% fetal bovine serum (Thermo Fisher), and 50µg/ml penicillin-streptomycin (media called D10). All *T. gondii* strains were propagated *in vitro* by serial passage on HFF monolayers (69). *In vitro* tachyzoite-to-bradyzoite conversion was induced by exposing parasite cultures to pH 8.2 (70, 71).

### Generation of transgenic T. *gondii* strains

Pru*Δku80Luc* ((42) herein termed Pru) was used as parental strain to generate all transgenic lines used in this study. The parental strain expresses the enhanced green fluorescent protein (EGFP) when converted in bradyzoite due to the integration in its genome of an EGFP cassette expressing the fluorescent marker under control of the LDH2 bradyzoite stage specific promoter, as previously described (72).TgAAT1 knockout and all tagging strains were generated using CRISPR/Cas9 technology as described in Rivera-Cuevas et al. (73). Guide RNAs (gRNA) used for genome manipulations are enlisted in Table S1. Schematic representations of the genomic manipulation strategy and PCR validation of all strain generated are in supplemental figures. Plasmids used to overexpress TgAAT proteins via transient transfection of Pru were generated by cloning TgAAT cDNAs in the pTub/CAT vector (74) by replacing CAT gene using NEBuilder® HiFi DNA Assembly technology (NEB E5520S). TgAAT cDNAs were amplified using forward primers containing the coding sequence for the c-myc tag to fuse the c-myc epitope at the N-terminus of the TgAAT proteins. Complementation of TgAAT1 (PΔ*aat1*:AAT1) was accomplished by integrating a plasmid carrying the TgAAT1 cDNA cloned downstream of 1,581 bp of TgAAT1 sequence upstream of the initiation codon to drive transcription of these sequences, followed by 312 bp of the 3’UTR of the TgSAG1 gene and the BLE selection cassette. The plasmid was integrated into the *T. gondii* genome upstream of the tubulin gene by introducing in the complement plasmid a 1,300 bp fragment derived from the tubulin locus and linearization using the *Pme*I to induce single crossover, as described in Kannan et al. (63). *T. gondii* transfections were performed as described previously (75). Correct integrations was confirmed by PCR analysis of single clones using a Phire tissue direct PCR Master kit (Thermo) (76).

### Immunoblotting and immunofluorescence

Immunoblotting and immunofluorescence were performed as described (73). Mouse anti-c-myc was diluted 1:1000 and 1:100 in WB and IFA, respectively. Anti-TgCPL was used at 1:200 in IFA (42).

### Tachyzoite plaque assay

Intracellular tachyzoites were mechanically liberated and purified following standard procedures. Three hundred tachyzoites were added to HFF monolayers grown in 6-well plates in triplicate or quadruplicate wells. Parasites were left undisturbed for 12 days at 37°C and 5% CO_2_. Plates were then stained with crystal violet and plaques were counted.

### Competition assay

HFF monolayers were infected with 10^6^ parasites composed of a mixture of either two strains: Pru and PΔ*aat1* or Pru and PΔ*aat1:AAT1*, in 1:4 ratio. Parasites were collected after 5 passages of monolayer lysis and their genomic DNAs were analyzed by qPCR using the primers Tub-F1 and DHFR-R1 (listed in Table S1) designed to amplify 119 bp of the DHFR selection cassette (absent in Pru but present in PΔ*aat1* and PΔ*aat1:AAT1*). Amplification of the tubulin gene (present in all parasites) was used to normalize the gDNA quantity.

### PMA/qPCR assay

This new protocol was developed by modifying the PMA viability assay applied to bacteria and other eukaryotes (48–57). Exploiting propidium monoazide molecule (PMA), it is possible to discriminate dead from live cells. PMA is not permeable to membranes and thus enters only dead cells because membranes are damaged. Inside dead cells, PMA binds genomic DNA (gDNA). Exposition of these samples to 470 nm blue light covalently cross-links PMA to gDNA. This irreversible binding renders the gDNA not amplifiable by PCR. So, only the gDNA of live parasites, not accessible to PMA binding, can be amplified from samples incubated with PMA and exposed to blue light. Variability due to loss of material during centrifugation steps was minimized by directly performing qPCR on aliquots of PMA-treated and non treated parasites after adding the DNARelease Additive from the Phire tissue direct PCR Master kit (Thermo Fisher Scientific) without any gDNA purification.

**Tachyzoites viability assay** was performed by mechanically liberating parasites from infected monolayers, purification of tachyzoites from cell debris by filtration though 3 µm filters and incubation of 2×10^5^ extracellular parasites in 39µl of D10 with or without 30µM PMAxx (Biotium) for 15 min in shaking and darkness. Incubation with PMAxx was followed by blue light exposition for 15 min using a Blue LED Device (PMA-Lite™ LED Photolysis Device Biotium #E90002). After blue light exposition, 1µl of DNARelease Additive was added to each sample followed by incubation at 22°C for 4 min and at 98°C for 2 min in a PCR machine. One μl of DNA from each sample was used to perform qPCR in 10μl total mix from POWERTRACK SYBR MM (Thermo) using 0.3µM multilocus primers Tx9 and Tx11 (Table S1). PCR running parameters were set as follows: 95°C for 15 s and 60°C for 30 s for 40 cycles. Viability was calculated as follow:

- dCt_sample_ = Ct_(PMAxx-treated sample)_ - Ct_(untreated sample)_
- fold change = 2^dCt^sample
- % of viability = 100/fold change

**Bradyzoites viability assay** was performed as follow: 10^2^ tachyzoites per well were used to infect HFF monolayers grown in 96-well plates. Twenty-four h post infection, tachyzoites were induced to convert into bradyzoites by replacing D10 media with alkaline media for 1 week or 2 weeks, refreshing the media daily. After conversion, infected monolayers were washed 3 times with HBSS and bradyzoites were released from *in vitro* cysts by adding 33.2μl of pre-warmed pepsin solution (0.026% Pepsin in 170mM NaCl and 60mM HCl, final concentration) and incubation at 37°C for 1 h (HFF monolayer is degraded by this treatment). Reactions were stopped by adding 1 volume (33.2μl) of 188mM Na_2_CO_3_. Bradyzoite viability was performed by transferring 33.2μl from each sample into 2 new 96-well PCR plates for PMA-treated and untreated samples and placing the 96-well plates on ice. PMA treatment was performed by adding 5.8μl of 200μM PMAxx to the 33.2µl of the PMA-treated samples, while the same volume of ddH_2_O was added to the untreated tubes. Samples were incubated for 15 min in shaking and darkness and then exposed to blue light for 15 min using the PMA-Blue LED Device. One μl of DNArelease Additive was added to each sample followed by incubation at 22°C for 4 min and 2 min at 98°C in a PCR machine. One μl of each sample was used in qPCR to assess viability as described above for tachyzoites.

### Invasion assay

Invasion assay was performed as already described in Possenti et al. (77) by differential IFA using a red-green assay (78) by labeling exclusively attached (extracellular) tachyzoites in non-permeabilized samples with anti-TgSAG1 and, after permeabilization with 0.2% Triton X-100, all parasites were stained with rabbit anti-GAP45 antibody. 300 microscopic fields/well at 100× magnification were examined. The data shown are representative of experiments performed in triplicate.

### Replication Assay

Freshly egressed parasites were used to infect HFF monolayers grown on coverslips in 6-well plates at a density of 10^6^ tachyzoites/well for 20 min. After 4 washes to remove uninvaded tachyzoites, the cultures were incubated for 24 h at 37°C and 5% CO_2_ prior to fixation with 4% PFA. Infected monolayers were permeabilized with 0.2% Triton X-100 and processed for IFA using rabbit anti-GAP45 and 594-conjugated anti-rabbit IgG antibodies. The number of parasites per vacuole was enumerated by examining 300 microscopic fields/well at 100× magnification. The data shown are representative of experiments performed in triplicate.

### *In vivo* cyst viability

CBA/J female mice (7 to 8 weeks old; Jackson Laboratories, Bar Harbor, ME) were used in this study. Mice were injected intraperitoneally (i.p.) with purified 10^5^ tachyzoites in 200 µl of 1× phosphate-buffered saline (PBS). Five weeks post-infection, mice were sacrificed according to university-approved protocols. Brains were harvested and homogenized in 1ml of ice-cold PBS via syringing through a 20-gauge needle. Cysts were enumerated in 2×100µl of brain homogenate by fluorescence microscopy, and the total brain cyst numbers were calculated. Cyst burden data were pooled from three independent experiments.

### Bradyzoites Time Course

Freshly egressed tachyzoites were used to infect 20 HFF monolayers grown on coverslips for each Pru, PΔ*aat1* and PΔ*aat1:AAT1* strain and converted into bradyzoites by alkaline induction 24 h post-infection. Two slides per strain were fixed every day from day 1 to day 10 post alkaline induction to generate two series of time course conversion. One series for each strain was stained with mouse anti-TgSAG1 antibody (1:100) followed by Alexa 594- conjugated anti-mouse IgG antibody (1:1,000) and the second series was stained with rabbit anti-TgBAG1 antibody (1:400) followed by Alexa 594-conjugated anti-rabbit IgG antibody (1:1,000). 500 cyst images were collected to measure fluorescent intensity arbitrary units (a.u.) of TgSAG1, TgBAG1 and GFP staining. Because parasite vacuole/cyst images even when negative for a specific marker have a background that slightly changes from image to image, we have subtracted from the signal value of each marker the value obtained as average of the signal intensity collected from 500 Pru negative vacuoles at day 1 for TgBAG1 and EGFP signals, when these markers are not expressed, and at day 10 for TgSAG1, selecting negative cysts (some TgSAG1 positive tachyzoite vacuole were still present even after 7-10 days of alkaline induction).

### Esterified Amino Acid Viability assay

Intracellular parasites of Pru, PΔ*aat1* and PΔ*aat1:AAT1* strains were mechanically liberated from infected HFF monolayers by syringing through 26G needles, centrifuge at 800*g* for 10 min at 4°C and then resuspended in 1.5ml of D10. 10^6^ tachyzoites of each sample were transferred to each well of a 96-well plate with V bottom in a volume of 200 µl for each esterified amino acid or control analysis. After centrifuging the 96-well plate, parasites were resuspended in RPMI media where the amino acid to be tested was replaced with the esterified amino acid and then incubated at 37°C with 5% CO_2_ for 1 h. The quantity of each esterified amino acid supplemented to the RPMI media was selected as the highest that was not toxic to the parental strain. Viability of all samples was analyzed using PMA/qPCR protocol described above.

### PLVAC size assay

Pru-*FYVE-mCherry* (Pru*-FYmC*), PΔ*aat1-FYVE-mCherry* (PΔ*aat1-FYmC*) and PΔ*aat1:AAT1-FYVE-mCherry* (PΔ*aat1:AAT1-FYmC*) parasites were collected using the same protocol for esterified amino acid viability assay. After incubation, extracellular parasites were settled on Cell-Tak (Fisher Scientific) coated slides for 30 min at 4°C, fixed in 4% formaldehyde, and stained with rabbit anti-mCherry antibody (1:500, Merk) followed by Alexa 594-conjugated anti-rabbit IgG antibody (1:1000). Three hundred images for each sample were acquired by focusing on the PLVAC signal on a Nikon TE2000-S Inverted Fluorescence Microscope equipped with a 100× oil objective and processed using ImageJ software. PLVAC size was measured by defining the area of anti-mCherry immunofluorescence.

### Statistics

Data were analyzed using GraphPad prism. For each data set, outliers were identified and removed using ROUT with a Q value of 0.1%. Data were then tested for normality and equal variance. If the data passed, one-way or two-way analysis of variance (ANOVA) were performed. If the data failed, a Mann-Whitney U test was performed.

## ACKNOWLEDGMENTS

This work was supported by a US National Institutes of Health grant (R01AI120607 to V.B.C. and M.D.C.) and by the University of Perugia Fondo Ricerca Di Base 2021 and 2022 program of the Department of Chemistry, Biology and Biotechnology (MDC).

## Figure legends

**Figure S1.**
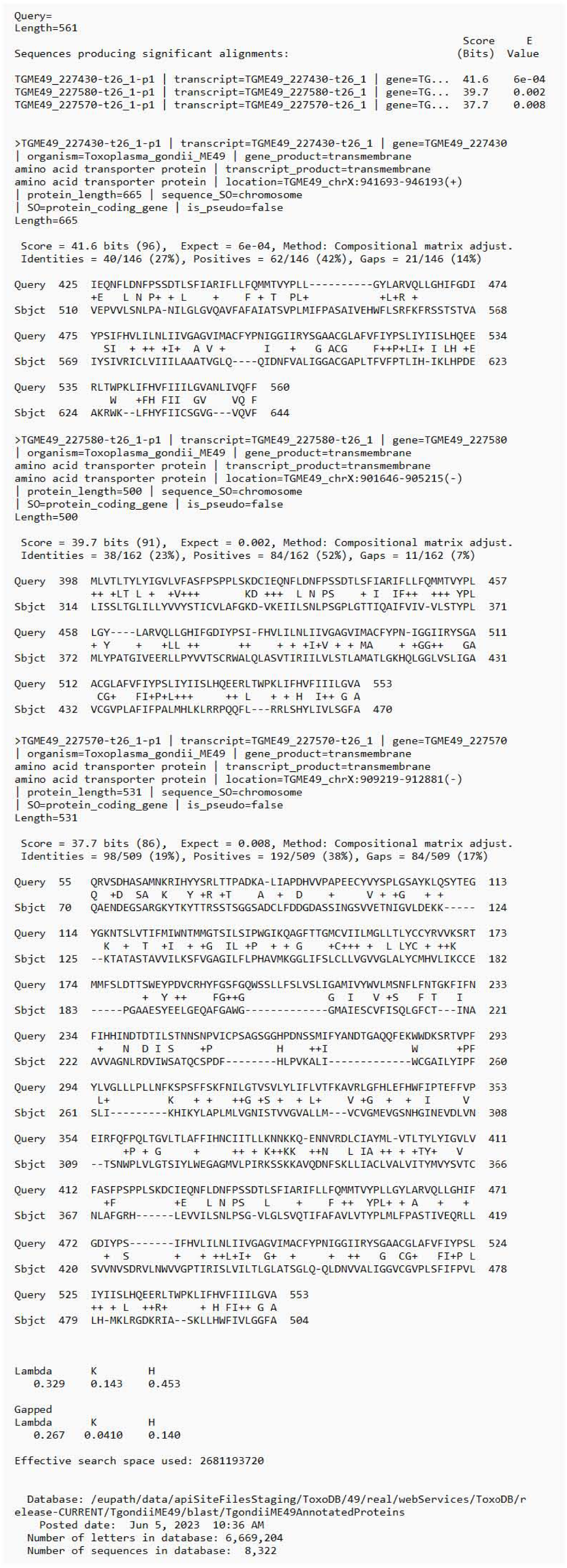
Search for putative SLC38A9 homologues in *T. gondii*. SLC38A9 was used as query to interrogate ToxoDB database (https://toxodb.org/toxo/app) to identify amino acid transporters potentially localized to the parasite PLVAC.

**Figure S2.**
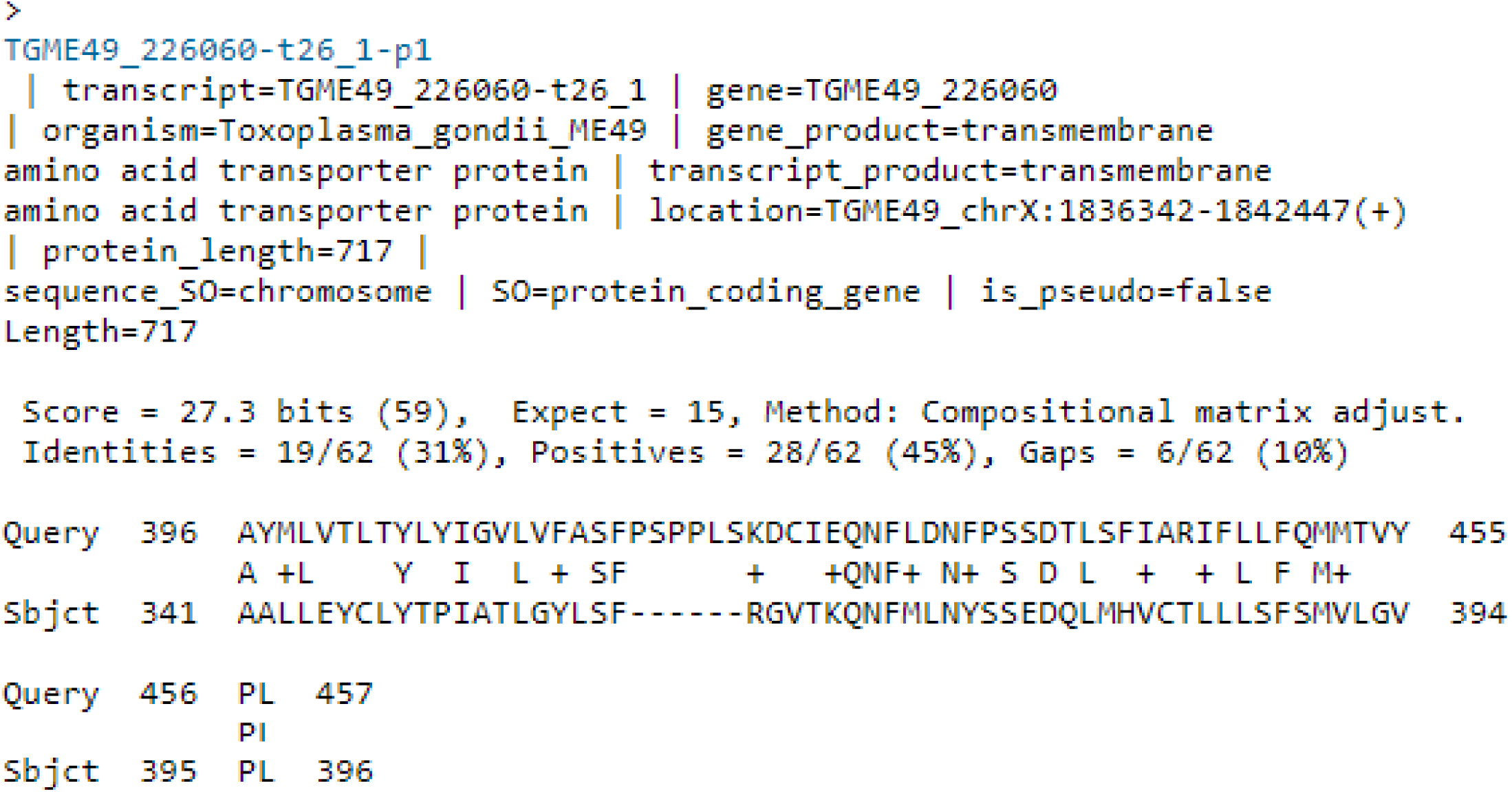
Identification of the fourth candidate as possible SLC38A9 homologue. Changing the setting of the expectation value in BLAST parameters from the default value of 10 to 20, the search for putative SLC38A9 homologues also identified the *T. gondii* protein TGME49_226060.

**Figure S3.**
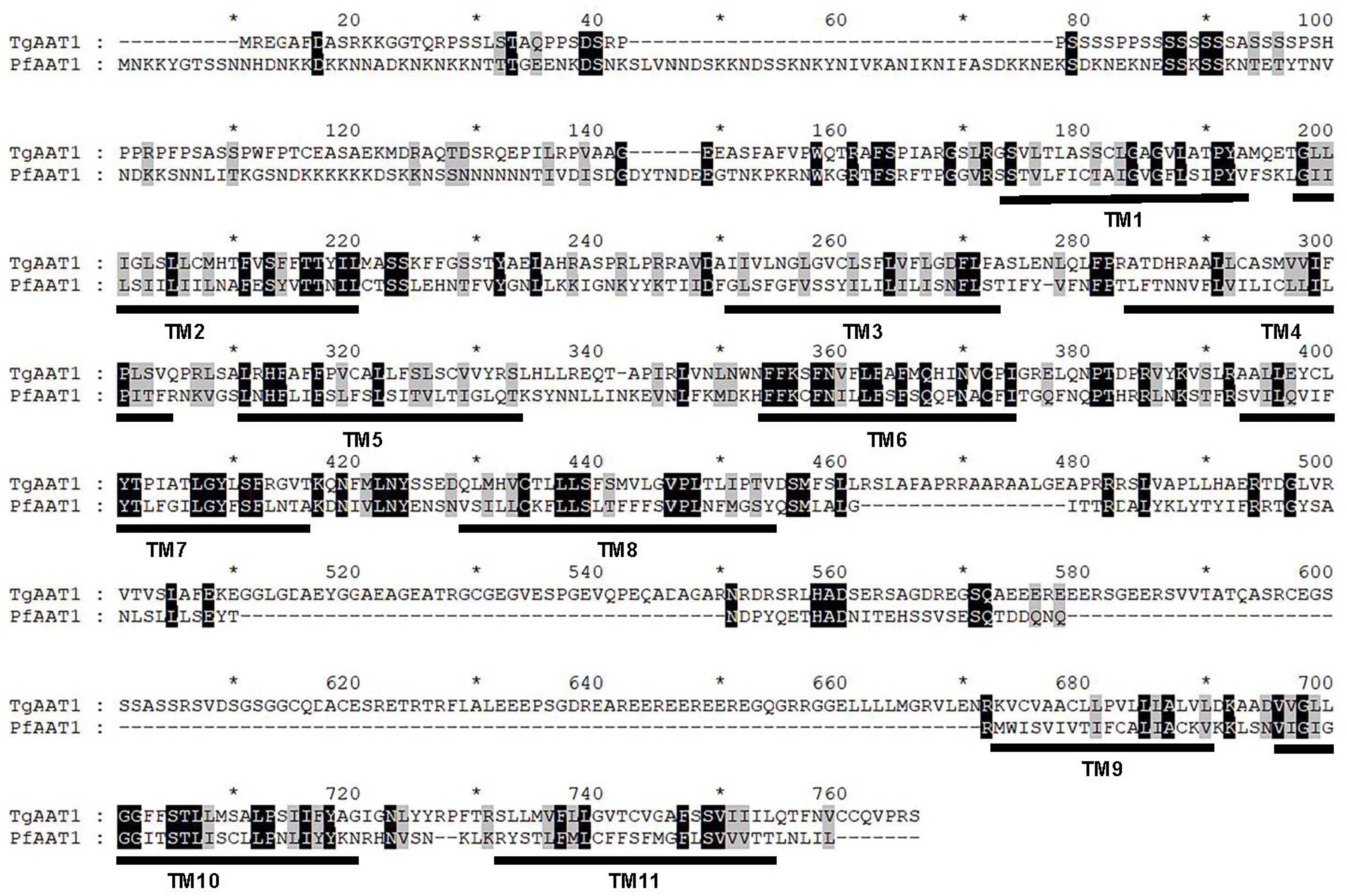
Sequence alignment between TgAAT1 and PfAAT1 using ClustalW software (https://www.genome.jp/tools-bin/clustalw). Transmembrane domains are underlined and number as TM1-11. PlasmoDB (https://plasmodb.org/plasmo/app) accession number of PfAAT1 is PF3D7_0629500.

**Figure S4.**
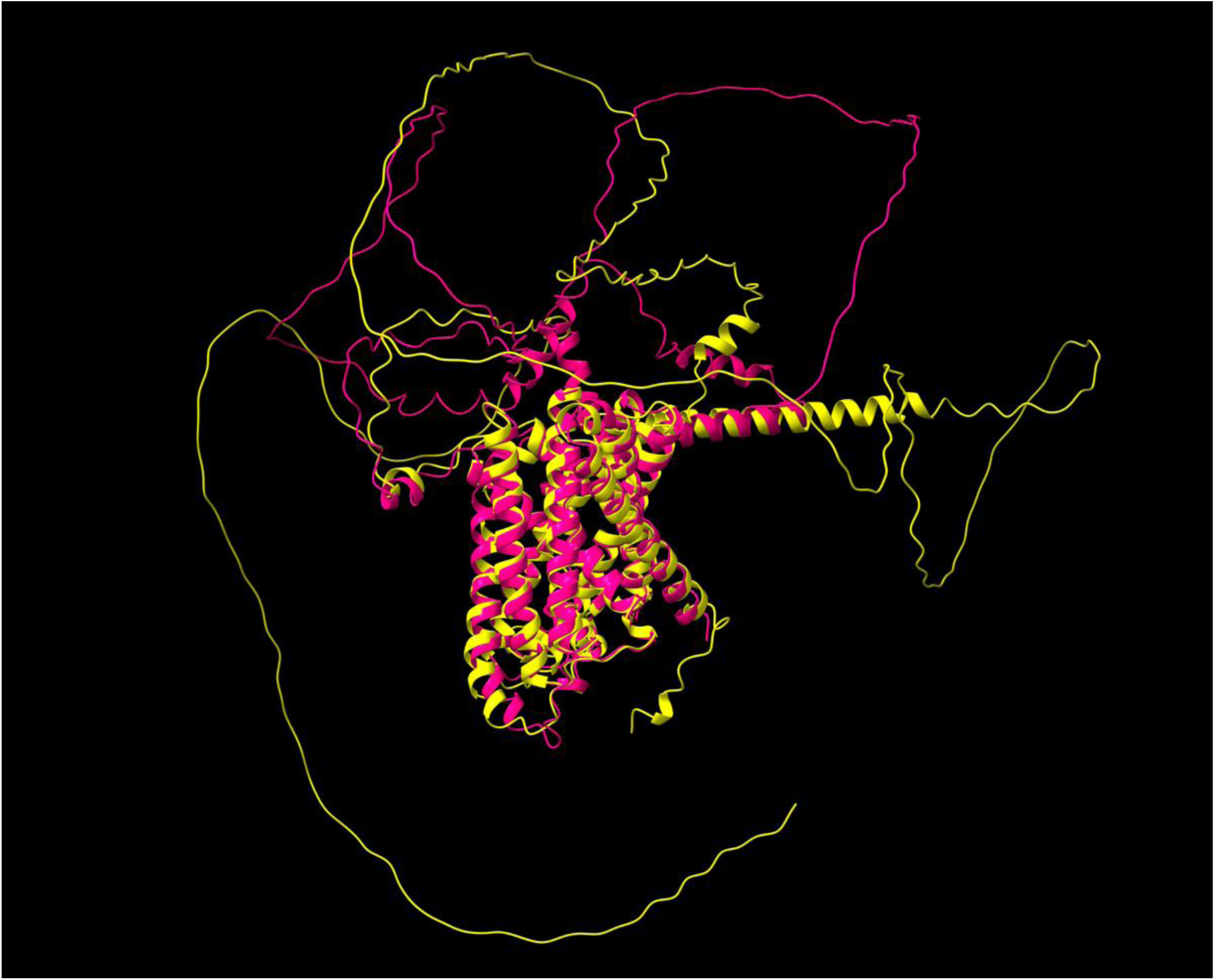
AlphaFold2 (https://alphafold.ebi.ac.uk/) model and structural comparison (http://ekhidna2.biocenter.helsinki.fi/dali/) of PfAAT1 (Pink) and TgAAT1 (Yellow).

**Figure S5.**
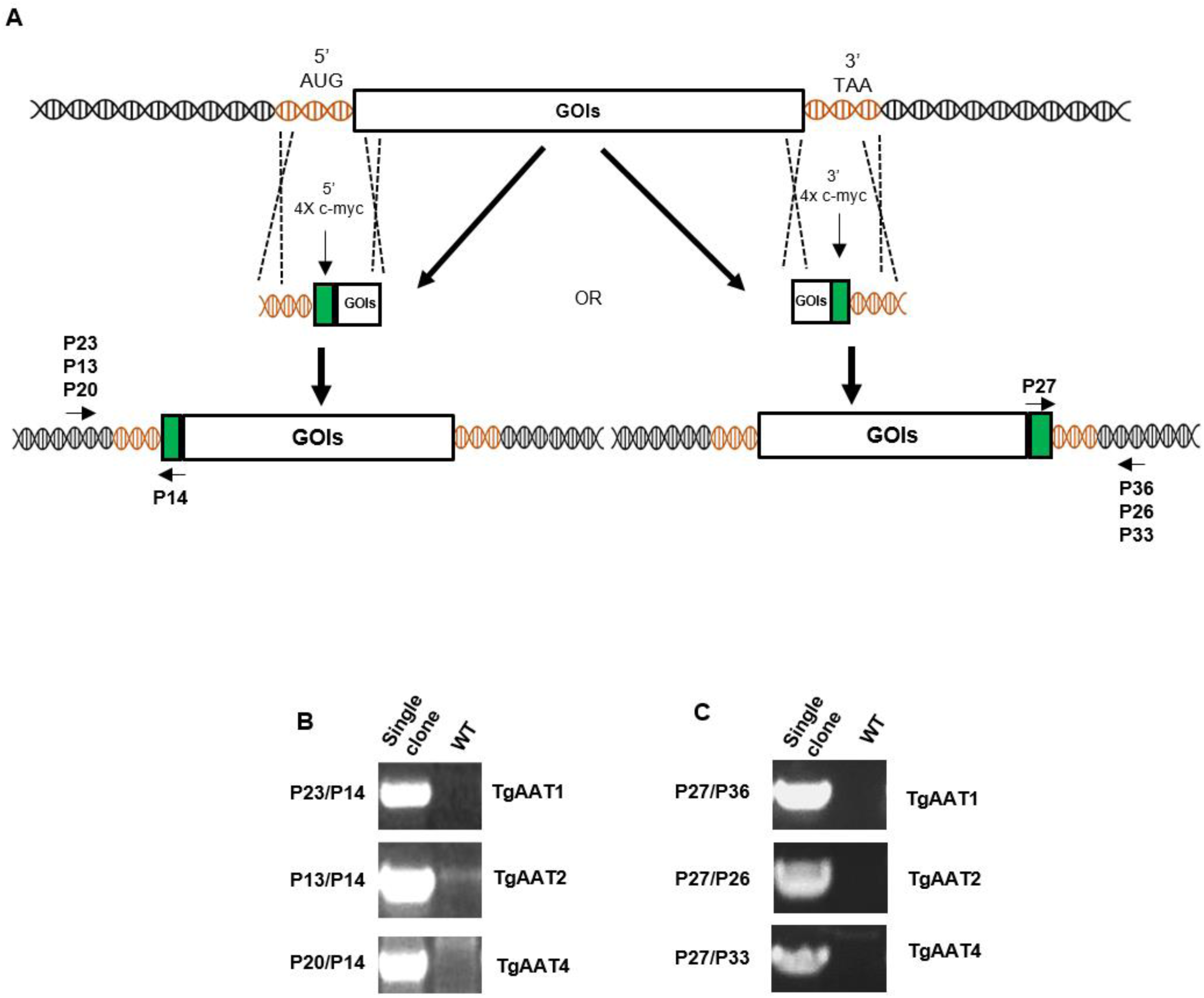
Strategy to endogenously tag TgAATs with 4x c-myc epitopes. A) A plasmid carrying the sequence encoding 4xc-myc epitopes was used to generate ⁓2µg of PCR fragment composed of the tag sequence flanked at both sides by 40 bp of homology to regions immediately adjacent to the sequences coding either the N-terminal (left panel) or C-terminal (right panel) regions of the TgAAT proteins. Integration of the 4x c-myc tag in the selected place by double-crossover homologous recombination was induced by Cas9 cleavage directed by a specific gRNA. Validation of correct epitope sequence integration was validated by PCR analysis. Panel B shows an agarose electrophoresis gel of the PCR screening products of single clones tagged at the N-terminus. The same analysis was employed for the C-terminus tagging (panel C). All primer sequences and gRNAs are reported in Table S1.

**Figure S6.**
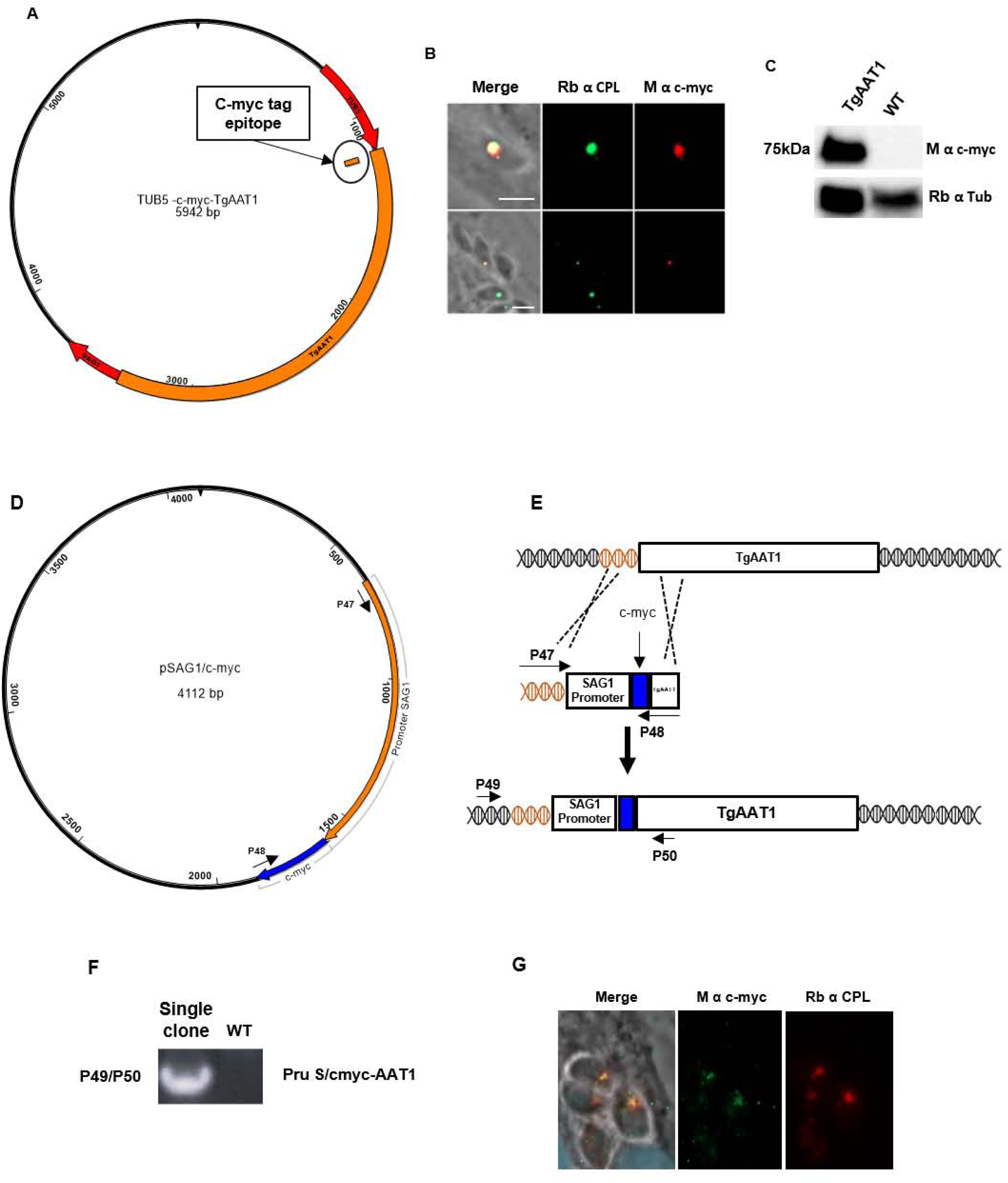
Panels A-C. Transient overexpression of TgAAT1. A)Map of the plasmid generated to overexpress TgAAT1 fused at its N-terminus to a single copy c-myc epitope. B) Representative images of parasites transiently transfected with the TgAAT1 overexpressing vector and processed for IFA using anti-c-myc (red) and anti-TgCPL (green) antibodies. Scale bar 10µm. C) WB analysis of the protein extract of parasites transiently transfected with the TgAAT1 overexpressing vector. The filter was first probed with rabbit anti-c-myc and then with anti-tubulin as a loading control. Panels D-G. Replacement of the TgAAT1 promoter with that of the TgSAG1 gene enabled the visualization of TgAAT1 through immunofluorescence assay (IFA). D) Map of the plasmid used to amplify the repair template consisting of the TgSAG1-c-myc cassette (TgSAG1 promoter transcribing the sequence encoding the c-myc epitope fused to the N-terminus of a gene of interest, GOI) flanked on both sides by 40 bp of homology to regions immediately adjacent to the TgAAT1 promoter. Primers and gRNAs used are reported in Table S1. E) Schematic representation of the strategy used to replace the TgAAT1 promoter with that of TgSAG1 gene. F) PCR validation of correct replacement of the TgAAT1 promoter with that of the TgSAG1 gene in a single clone. G) Representative images of parasites expressing TgAAT1 under the TgSAG1 promoter processed for IFA using anti-c-myc (red) and anti-TgCPL (green) antibodies. Scale bar 5µm.

**Figure S7.**
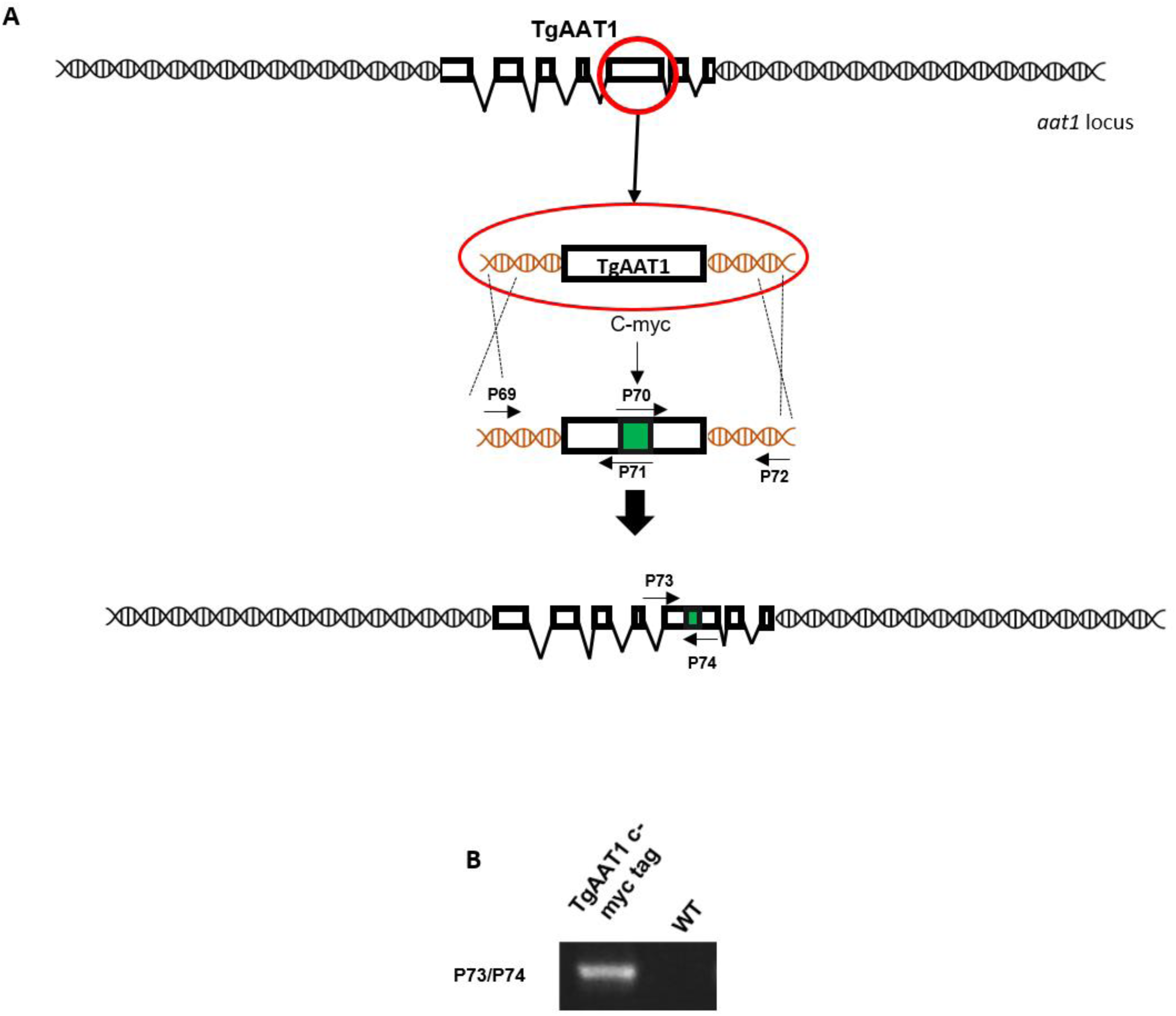
C-myc internal tagging of TgAAT1. A) Schematic representation of the strategy to internally tag TgAAT1. Overlap PCR were employed to generate a repair template consisting of the sequence encoding the c-myc epitope flanked at both sides by homology to regions immediately adjacent to the nucleotide triplet encoding the residue 508 of the TgAAT1 protein. Primers P69/P70 and P71/P72 were used to generate the two PCR products that were fused by overlap PCR using primers P69/P72 to create the repair template. B) PCR validation to confirm the successful insertion of the c-myc coding sequence in the intended place within the TgAAT1 gene in the Pru strain. All primer sequences and gRNAs are reported in Table S1.

**Figure S8.**
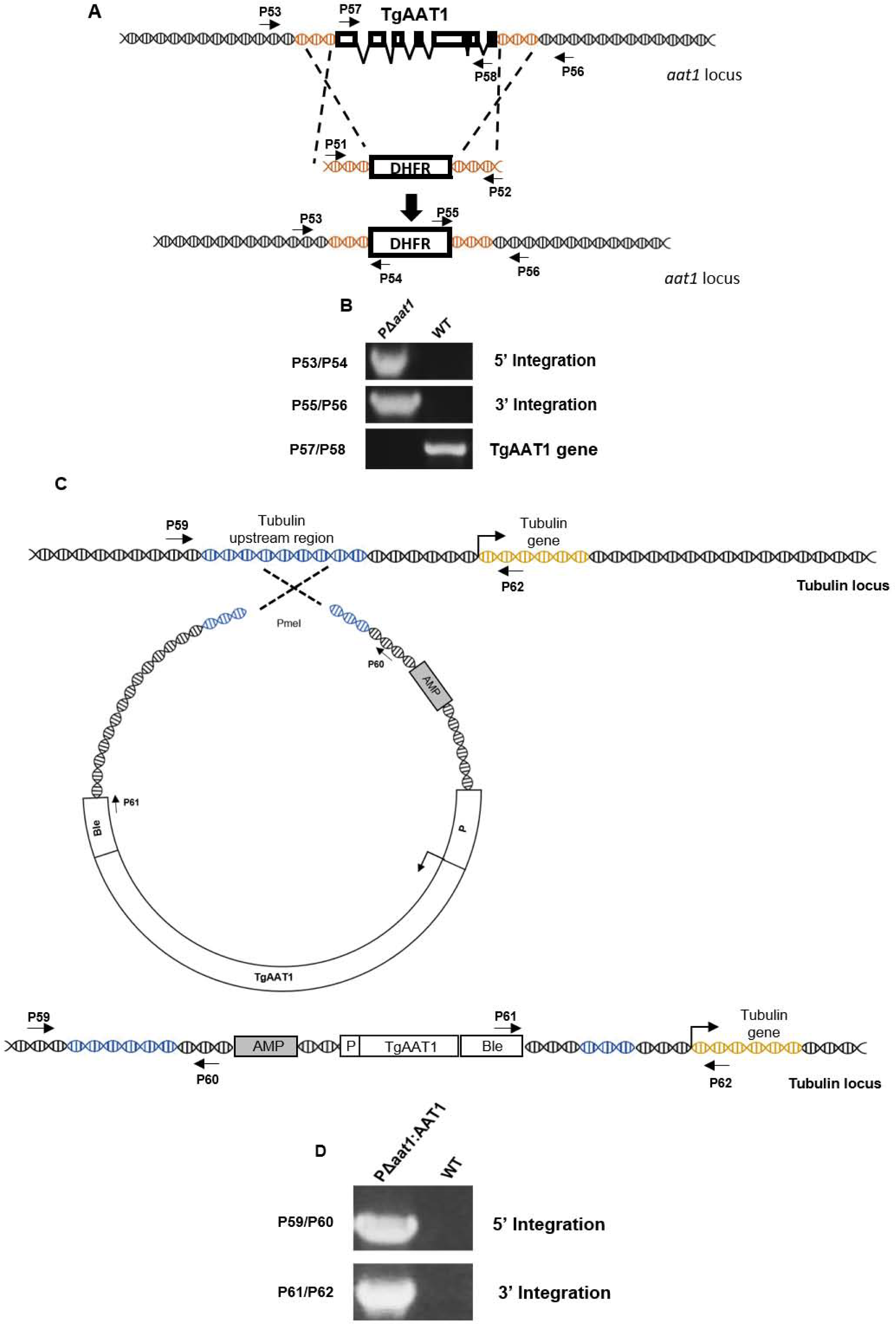
Panels A-B. Generation of the PΔ*aat1* strain. A) Schematic representation of the strategy used to delete the TgAAT1 gene. To generate TgAAT1 knockout parasites, two different gRNAs were used for Cas9 to cleave at both the start and stop codon *TgAAT1*. A repair template composed of the DHFR selection cassette flanked at both sides by 40 bp of homology to regions immediately adjacent to the start and stop codon of the TgAAT1 gene was used to select knockout parasites resistant to pyrimethamine selection. B) PCR validation to confirm the successful deletion of the TgAAT1 gene in the generated knockout parasites. Panels C-D. Creation and validation of genetically PΔ*aat1*:AAT1 complemented parasites. C) Schematics of single crossover integration to restore expression of TgAAT1 via genetic complementation with constructs containing a bleomycin selection cassette (BLEO). Single crossover integration was directed to the tubulin locus via homologous regions indicated in blue. Primers used to assess integration of the complementation construct are shown as numbered arrows. D) PCR analysis of integration at the tubulin locus using primers shown in C). All primer sequences and gRNAs are reported in Table S1.

**Figure S9.**
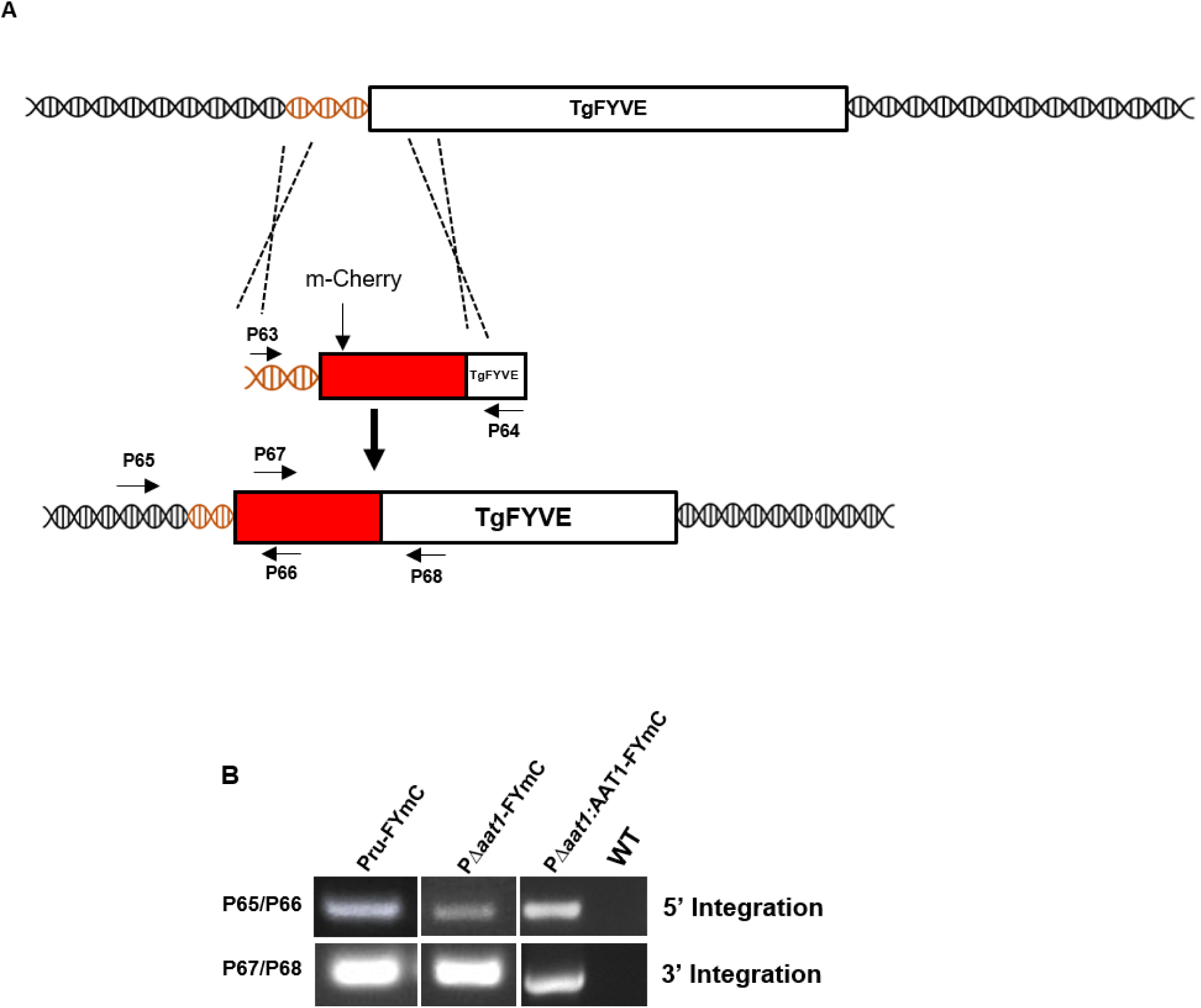
N-terminus tagging of TgFYVE with mCherry in Pru, PΔ*aat1* and PΔ*aat1:AAT1*. A) Schematic representation of the strategy to tag TgFYVE with mCherry. The repair template used to generate this strain contained the mCherry coding sequence flanked by 40 bp homology to the sequence upstream and downstream the start codon of the TgFYVE gene. Integration of the repair template in the intended place was achieved co-transfecting parasites with a plasmid expressing Cas9 and a gRNA that guided the nuclease to cleave close the start codon of the TgFYVE gene. B) PCR validation to confirm the successful integration of the mCherry coding sequence at the start codon of the TgFYVE gene in Pru, PΔ*aat1* and PΔ*aat1:AAT1* parasites. All primer sequences and gRNAs are reported in Table S1.

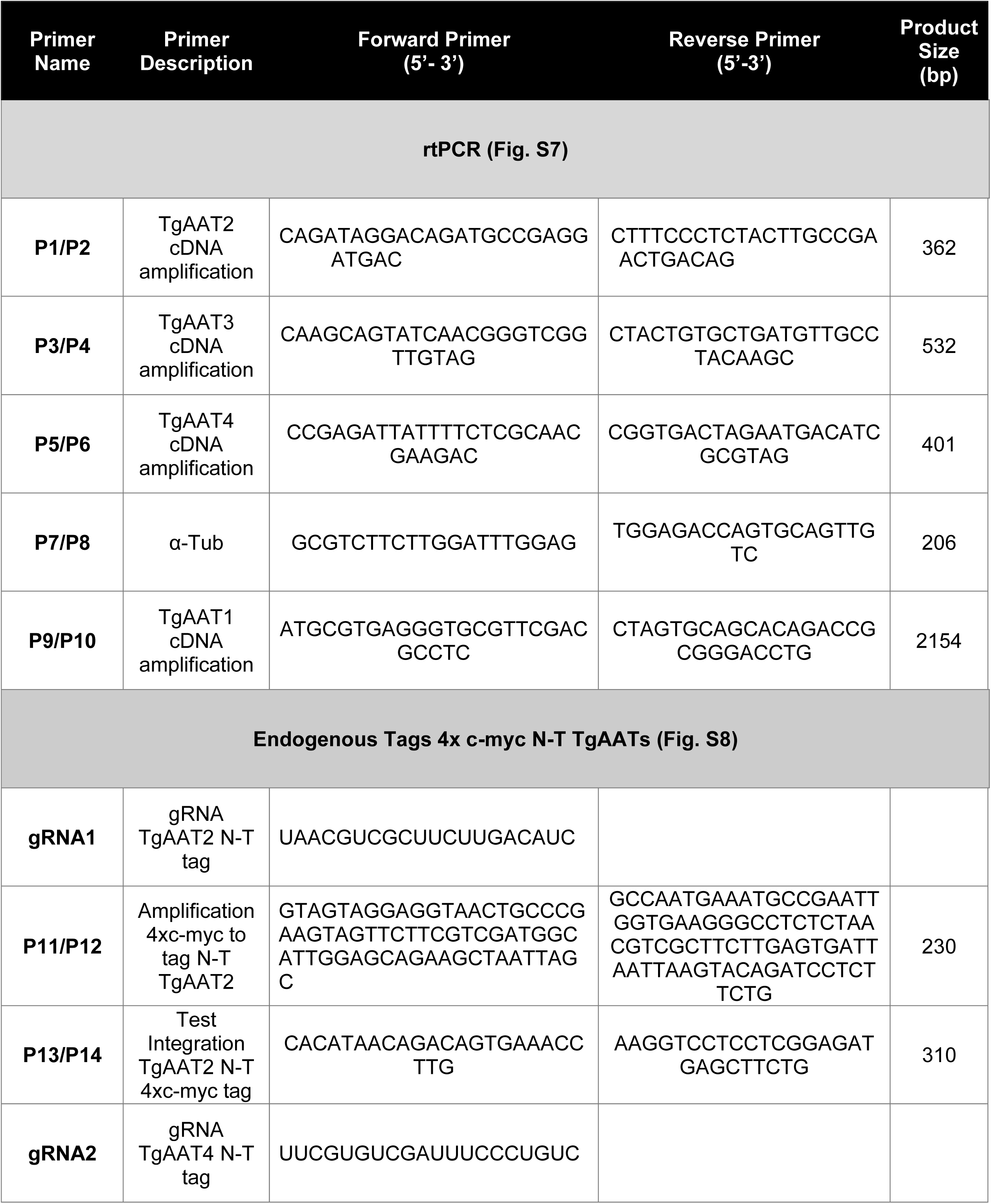

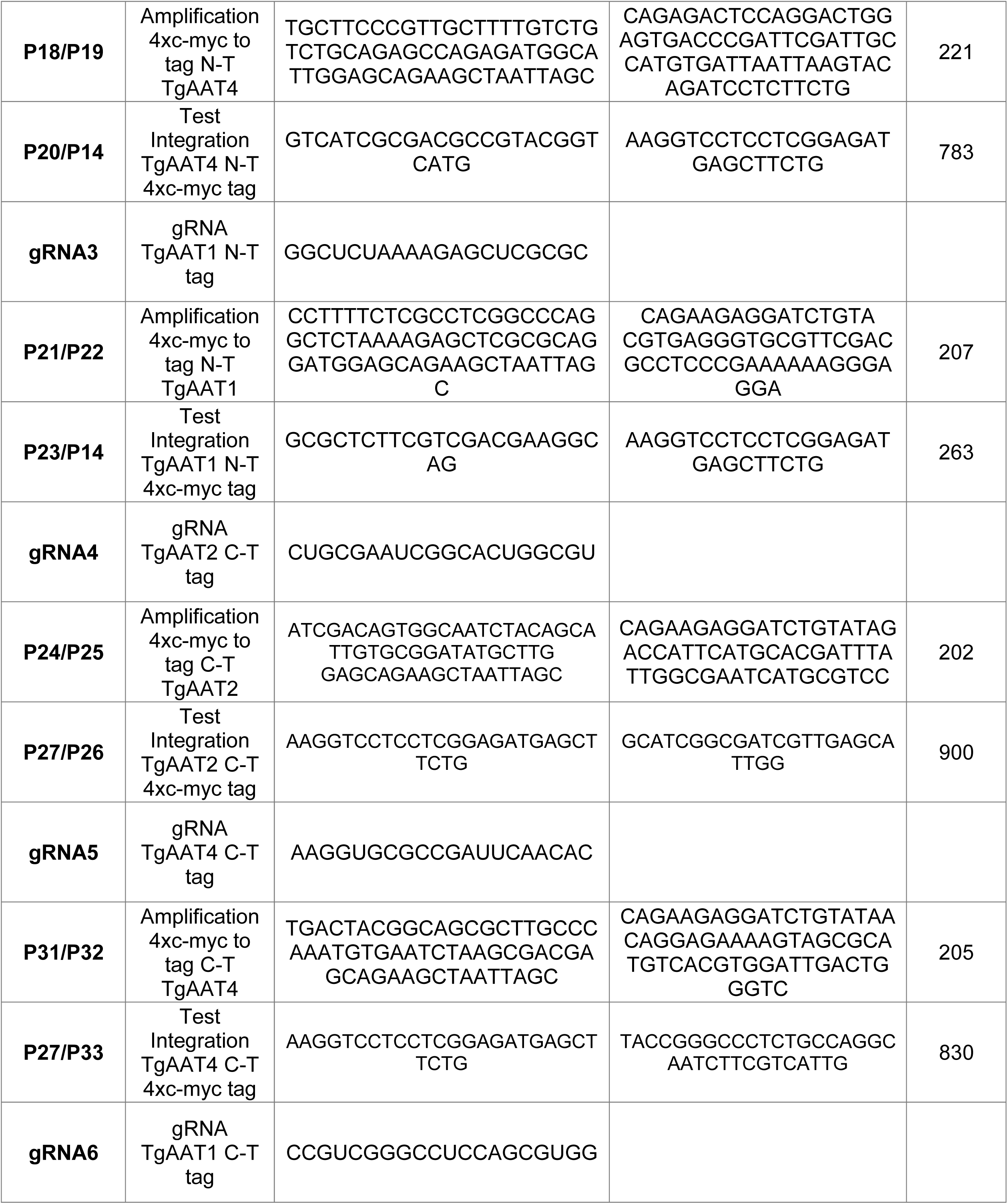

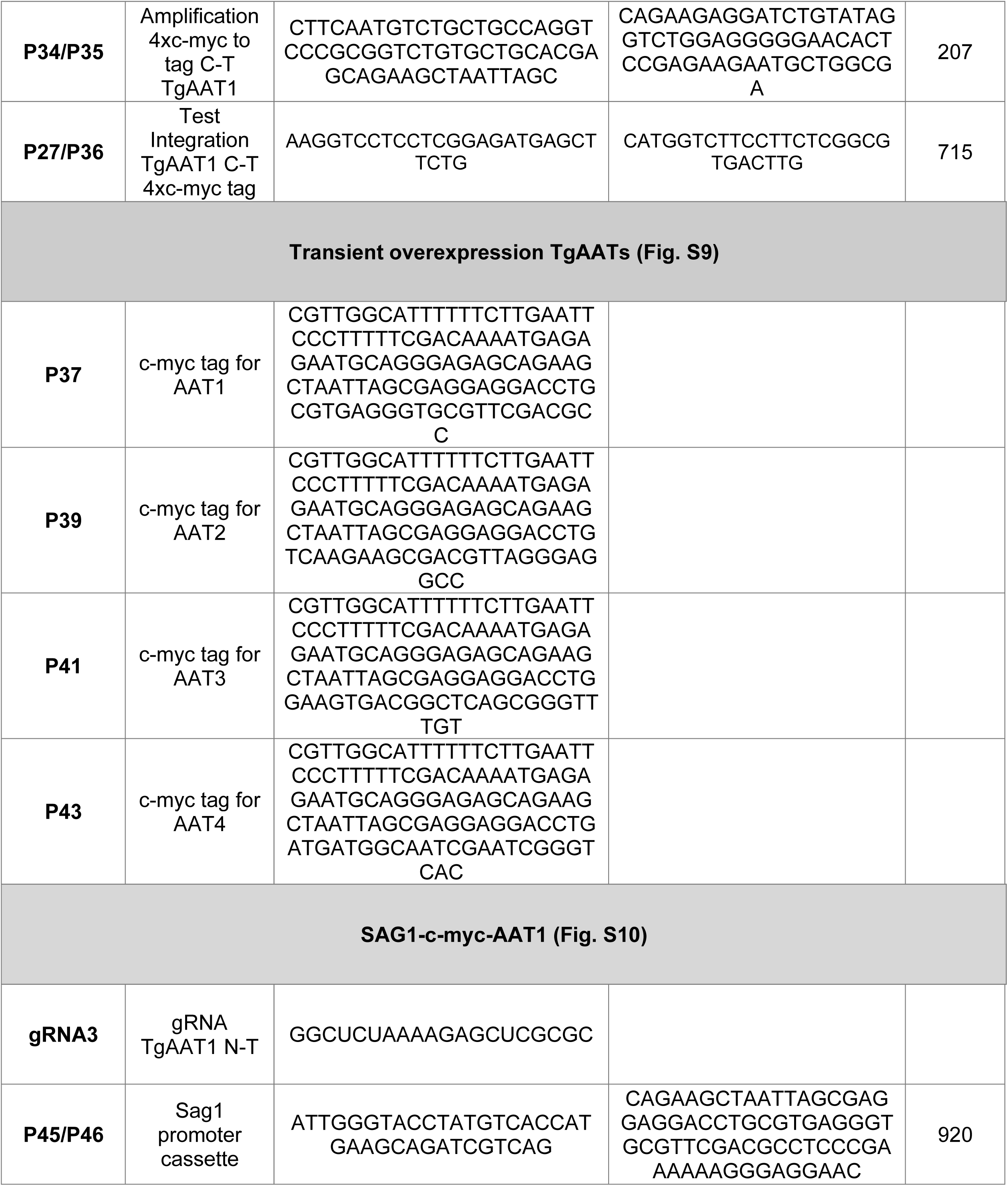

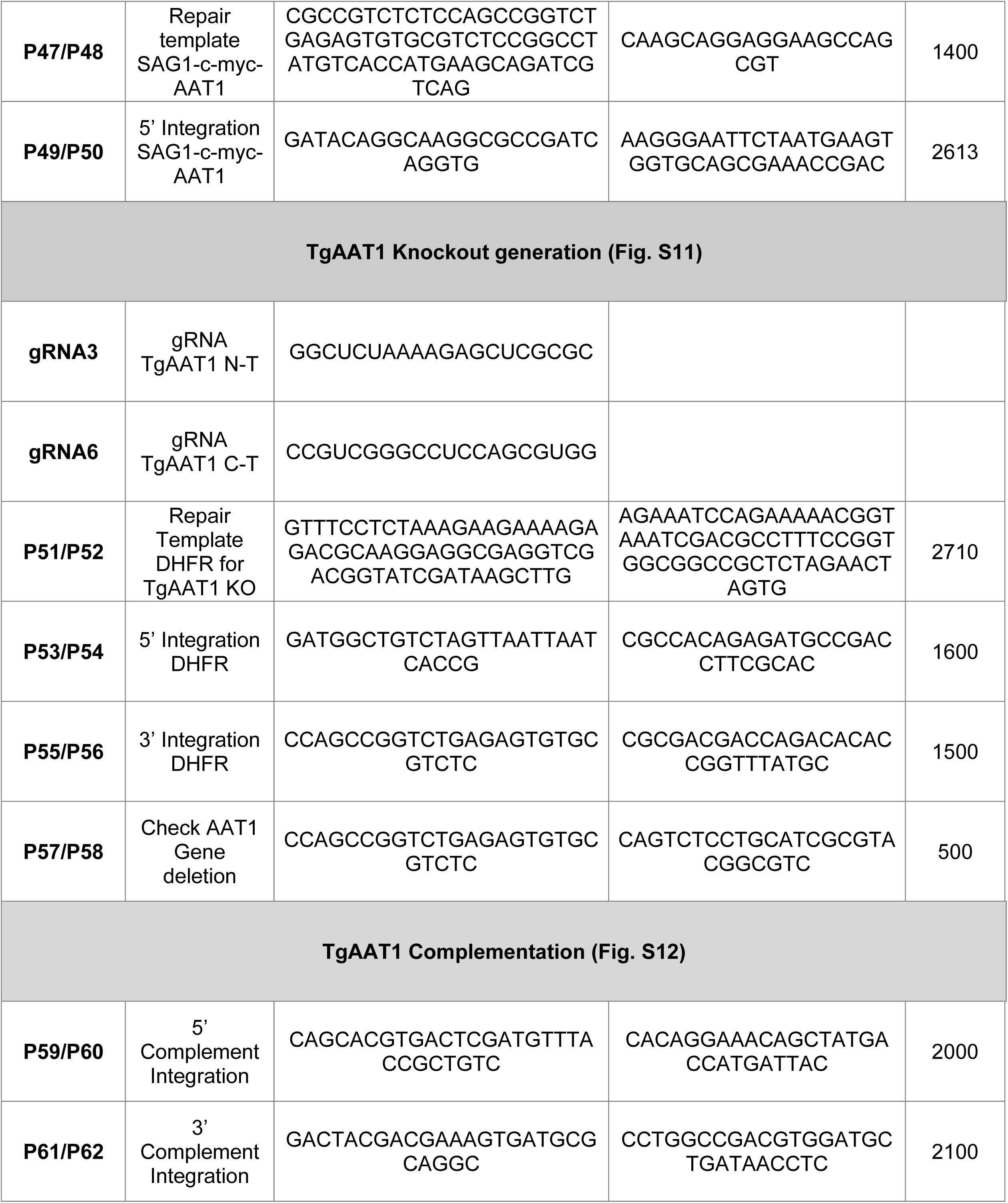

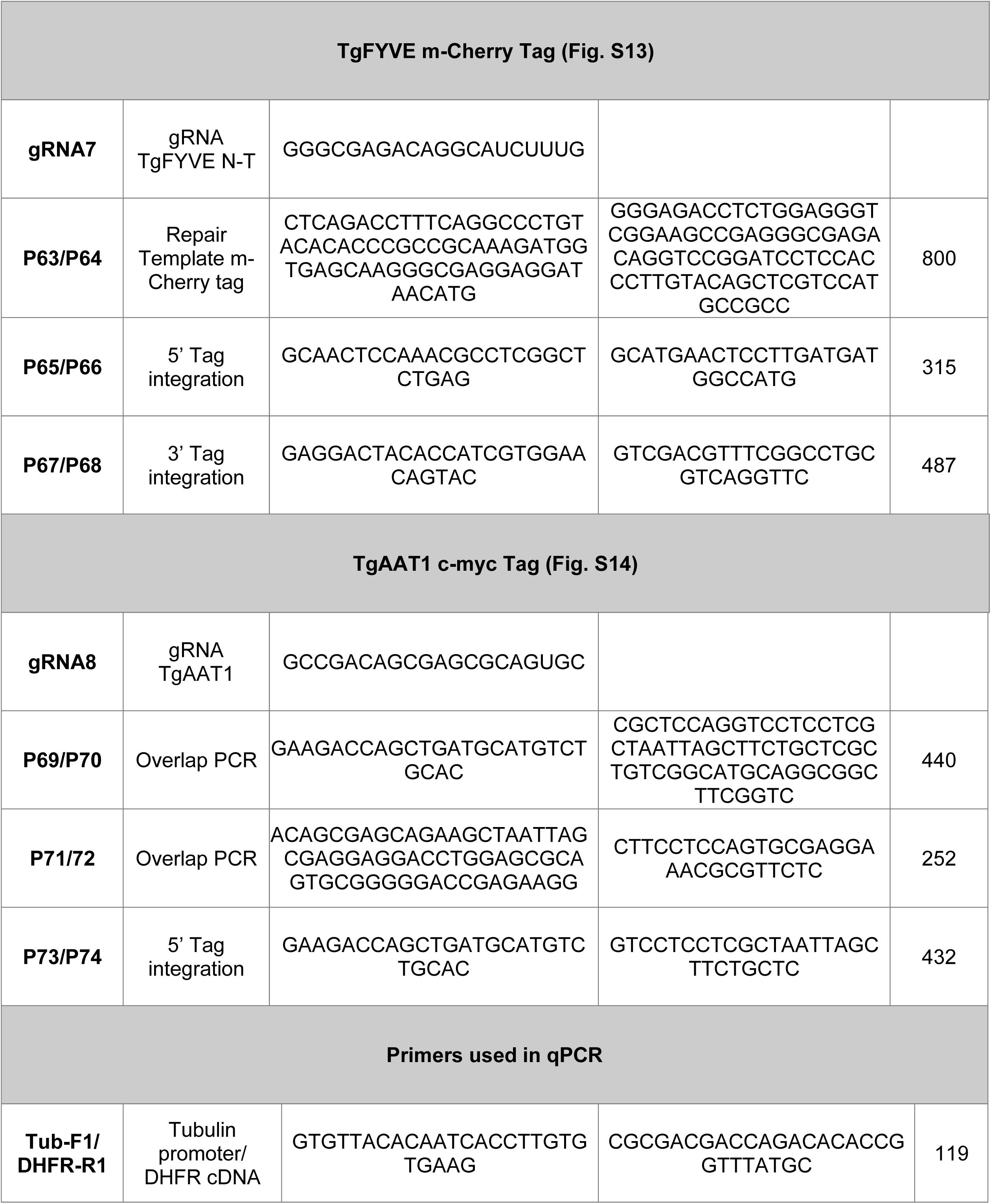

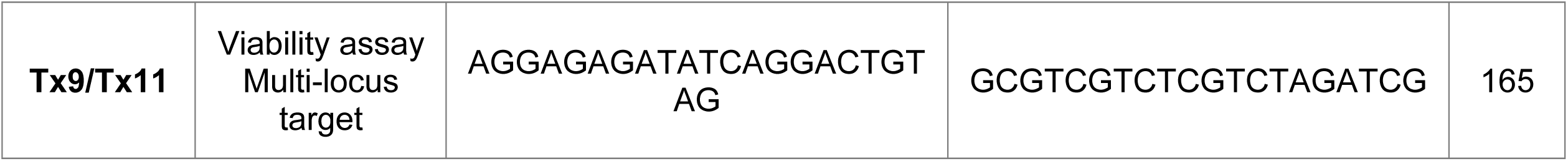

